# *De novo* Golgi biogenesis requires coordinated transactivation of a Golgi regulon

**DOI:** 10.1101/2025.09.17.676727

**Authors:** Francesca Forno, Domenico Abete, Elena V. Polishchuk, Xabier Bujanda Cundin, Fioranna Renda, Roberta Crispino, Josephine Salzano, Raffaella Petruzzelli, Rossella De Cegli, Martina Sofia, Nicolina Cristina Sorrentino, Lorenzo Vaccaro, Davide Cacchiarelli, Juul Verbakel, Jan De Boer, Bruno Goud, Alexey Khodjakov, Franck Perez, Roman S. Polishchuk

**Affiliations:** Telethon Institute of Genetics and Medicine, Pozzuoli (NA), Italy; Institut Curie, Paris, France; Wadsworth Center, New York State Department of Health, Albany (NY), USA; Eindhoven University of Technology, Eindhoven, The Netherlands

## Abstract

The Golgi apparatus expands during differentiation and high secretory demand, yet the transcriptional control of its biogenesis remains poorly defined. Here, we developed a targeted enzymatic ablation method to eliminate the Golgi and trigger *de novo* organelle formation. Single-cell RNA-seq of cells rebuilding Golgi revealed an orchestrated induction of a broad Golgi gene network coinciding with structural and functional organelle maturation. This gene set spans all Golgi sub-compartments and functions, including glycosylation, trafficking, and ion transport, thus supporting the concept of a unified Golgi regulon, enabling the simultaneous expression of components required for the organelle structural and functional integrity. Through promoter analysis and RNAi screening, we identified CREB3L1 as a key transcriptional regulator critical for Golgi gene activation and organelle reformation. These findings indicate that CREB3L1-dependent transcriptional mechanisms orchestrate a complete Golgi biogenesis program that may be essential for organelle regeneration and for secretory pathway plasticity during physiological remodeling.

## INTRODUCTION

Biogenesis of the Golgi apparatus is a vital process that occurs in various physiological contexts, particularly during increased secretory demand and cell differentiation. Golgi expansion is observed in various secretory cell types, including pancreatic beta cells, thyroid cells, and mammary acinar cells, upon stimulation ^1–3^. In B cell differentiation, for example, the Golgi volume increases nearly sevenfold to support antibody secretion ^4^. New Golgi units emerge in differentiating neurons and muscle cells ^5,6^, and in organisms such as *Giardia lamblia*, Golgi-like structures appear only at specific life stages ^7^, underscoring the organelle dynamic and responsive nature.

Golgi biogenesis requires the synthesis of proteins, lipids and other macromolecules to support organelle growth and function. However, the transcriptional mechanisms coordinating these biosynthetic processes with Golgi morphogenesis remain incompletely understood. A major barrier to dissecting this regulation has been the absence of an experimental system capable of triggering synchronized, large-scale formation of new Golgi structures from a depleted state. Such a system would ideally initiate from near-complete Golgi loss, requiring cells to regenerate morphologically and functionally mature Golgi stacks using newly synthesized components.

Complete Golgi disassembly can be achieved by treating cells with Brefeldin A (BFA), but upon drug removal, the Golgi rapidly reassembles from preexisting components without requiring new protein or lipid synthesis ^8^. Because this bypasses *de novo* biosynthesis, BFA washout may not fully capture the requirements of *de novo* Golgi biogenesis.

To address this, alternative models have been explored. In our earlier work, we employed diaminobenzidine (DAB) to inactivate Golgi in mannosidase II-HRP (ManII-HRP) expressing and, thereby, initiate new Golgi formation. Although Golgi-like membranes appeared within 2–6 hours, they failed to form stacked cisternae or support cargo trafficking ^9^. Moreover, Golgi inactivation has been shown to impair mitotic progression and promote cell death ^10^.

Another strategy—laser-mediated removal of the Golgi-containing region—elicits organelle regeneration in the remaining portion of the cell ^11,12^. Notably, this approach revealed that Golgi regeneration occurs slowly, with discernible stacks forming only after ~12 hours. However, its technical complexity and low throughput limit its broader application.

The slow kinetics of Golgi biogenesis prompted us to revisit the enzyme-inactivation method with extended recovery time to enable full regeneration of functional Golgi. Here, we demonstrate that selective inactivation of the preexisting Golgi triggers the formation of new, morphologically and functionally mature Golgi structures in a significant fraction of cells. This system provides a unique platform to investigate *de novo* Golgi biogenesis. Using single-cell RNA sequencing, we identify a robust transcriptional program activated during this process, including the upregulation of a broad Golgi gene network, implicating transcriptional regulation as a critical component of Golgi biogenesis.

## RESULTS

### Cell recover Golgi stacks after enzyme-mediated inactivation of preexisting organelle

We observed before that DAB-based inactivation of Golgi structures in HeLa cells expressing ManII-HRP induced the formation of a “new”, but immature, Golgi-like structures which fail to support trafficking ^9^. To test whether fully functional Golgi might be recovered, we first optimized HRP-mediated inactivation procedure to improve cell survival (see Methods). Second, we extended the observation to longer time intervals (up to 24h) that may be required to complete *de novo* Golgi biogenesis ^11,12^ and investigated the formation of new Golgi structures by electron microscopy (EM). We found that immediately after treating the cells with low dose of H_2_O_2_, precipitated DAB filled all the cisternae in most Golgi stacks (Fig. 1A, Fig. S1A). In a small subset of the stacks (less than 10%) only the cis-most or trans-most cisternae remained DAB-negative. The other intracellular structures, including the Golgi-adjacent ER, ER exit sites, endosomes, and lysosomes (Fig. 1A), were not positive for DAB. This indicates that HRP-mediated inactivation specifically targets and effectively eliminates the pre-existing Golgi, without affecting other organelles or compartments.

**Figure 1.**
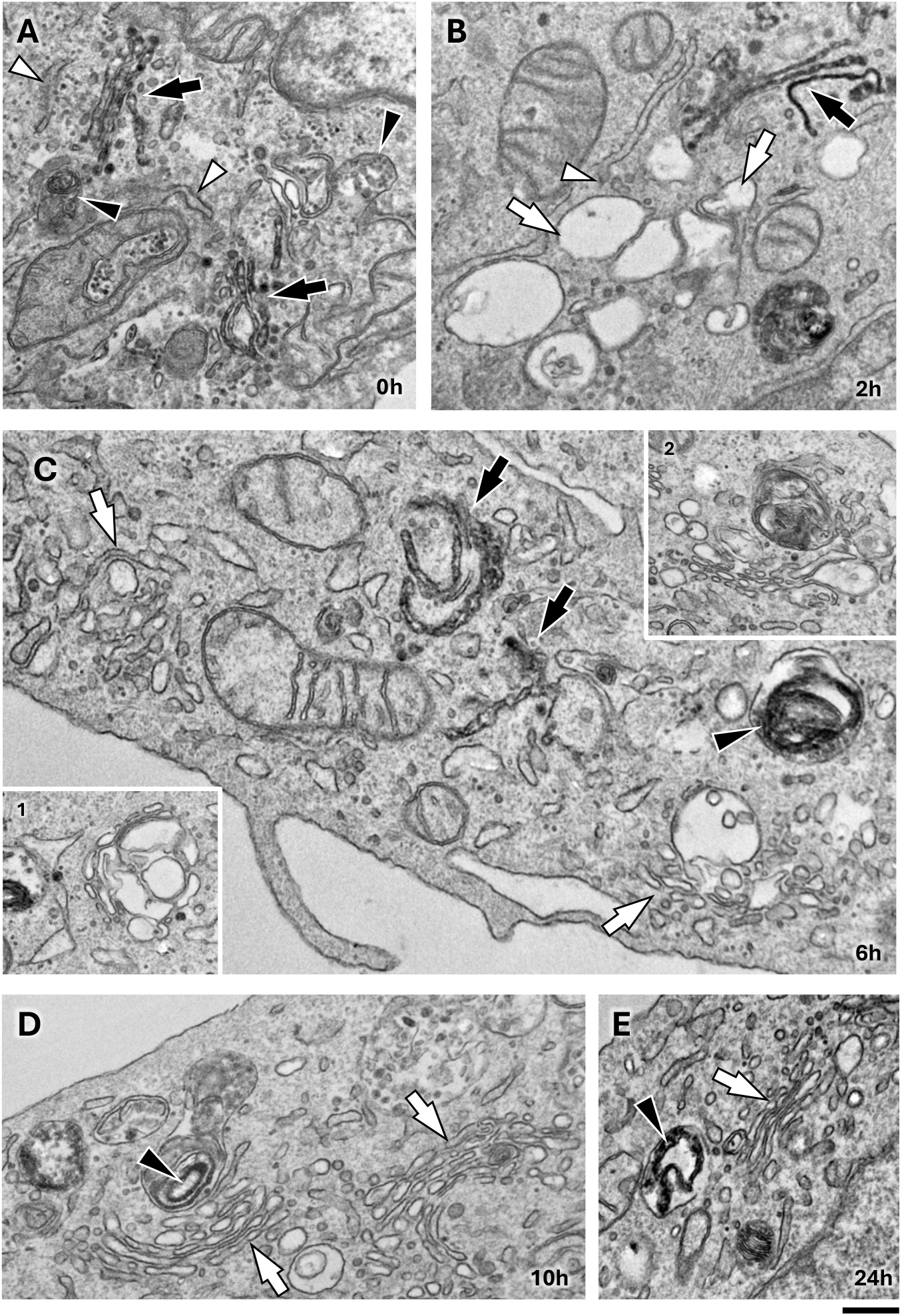
HeLa cells recover Golgi stacks after inactivation of preexisting Golgi organelle. ManII-HRP-expressing HeLa cells were incubated with DAB and H_2_O_2_ to cross-link the preexisting (old) Golgi and fixed for EM immediately (A) or after incubation with fresh medium for 2 (B), 6 (C), 10 (D) or 24 (E) hours. **A**. Black arrows indicate Golgi stacks with electron dense DAB polymer, which was deposited within cisternae across entire stack. Endolysosomal structures (black arrowheads) and ER cisternae (white arrowheads) did not contain DAB precipitate. **B-E**. DAB-negative newly-forming Golgi structures (“new” Golgi) are indicated by white arrows and were represented initially by vacuolar clusters (B) located near ER exit sites (white arrowhead in B), irregular stacks (C and insets 1, 2) or by regular stacks (D, E). Remnants of DAB-positive old Golgi were detected in cytosol (black arrows) or sequestered inside lysosomes (black arrowheads). Scale bar. 360 nm (A-E).

Two hours after inactivation, we observed clusters of vacuolar membrane-bound structures located near the old DAB-positive Golgi remnants (Fig. 1B; Fig. S1B). Frequently, the membranes within these clusters appeared stacked together (Fig. 1B, white arrows), and some began to flatten, acquiring cisternae-like shapes (Fig. S1B, white arrow in panel 2). The ultrastructure of these membrane clusters resembled previously reported morphologies of newly forming Golgi units in the cells, whose old Golgi has been eliminated by either enzyme inactivation ^9^ or laser ablation ^11,12^. Further immuno-gold labeling for GM130 confirmed that these newly-forming clusters represented Golgi-like compartments (Fig. S2A).

We also noted that these Golgi-like structures were often associated with ER exit sites (ERES) (Fig. 1B and Fig. S1B panel 1, white arrowheads), suggesting that their formation might be linked to the export of material from the ER. Notably, a few of these structures resembled Golgi stacks with markedly swollen cisternae, which we termed “irregular” stacks (or “irregular” Golgi) (Fig. S1B panel 3). The proportion of such irregular stacks significantly increased within the cell population by 6 hours post-inactivation of the old Golgi (Fig.1C, white arrows; Fig. S1C panels 1-3) and few Golgi structures resembling regular stacks were detected (Fig. 1C, inset 2).

By 10 hours, newly formed Golgi membranes began to adopt the stacked architecture but with flattened cisternae being observed in a substantial number of cells (Fig. 1D; Fig. S1D). Notably, the polarity of these newly formed Golgi stacks was restored: GM130 labeling was restricted to the *cis* side of the stack (Fig. S2B, white arrows), while clathrin-coated profiles were detected on the *trans* side (Fig. S2B, white arrowhead). Over time, the number of stacked structures increased in cells undergoing Golgi recovery (Fig. 1E; Fig. S1E, F), while vacuolar clusters and irregular stacks became less frequent, although they remained present in some cells (see quantification in Fig. S1F).

Importantly, we observed different phenotypes of newly forming Golgi within the same cell. During the early stages of Golgi recovery, vacuolar clusters were detected alongside irregular stacks (Fig. S2C). At later stages, such irregular stacks were seen coexisting with normal Golgi stacks within the same cell (Fig. S2D), suggesting that these phenotypes represent different stages of organelle biogenesis rather than distinct pathways of Golgi recovery.

We also observed that the old Golgi stacks underwent gradual disruption, with DAB-positive cisternae detaching from one another (Fig. 1B, C and S3A; black arrows). These DAB-positive cisternae were also found within autophagosomes (Fig. S3A, B; empty arrowheads) and within lysosome-like structures (Fig. 1C-E and Fig. S3A; black arrowheads). In agreement with previous study ^13^, we observed that, with the time significant portions of the old Golgi became decorated with the autophagic marker LC3 (Fig. S3C, D). Moreover, fragments of the old Golgi were also detected inside LAMP1-positive structures (Fig. S3E, F). This is consistent with EM observations showing the sequestration of DAB-positive cisternae within autophagosomes and autolysosomes. Apart from the appearance of autophagic and lysosomal structures sequestering the old Golgi, other intracellular organelles (such as mitochondria or the ER) appeared largely normal during Golgi recovery (Fig. 1). This indicates that the impact of Golgi inactivation is relatively specific and does not significantly affect other intracellular compartments or structures.

In sum, our EM observations indicate that cells are capable of rebuilding a new Golgi organelle with stacked architecture after the elimination of the pre-existing Golgi. However, this process requires significant time (usually ≥10 hours) and can vary considerably from one cell to another.

These observations were further confirmed by live cell imaging in ManII-HRP cells expressing the fluorescent Golgi marker GalT-GFP. We found that DAB deposition in preexisting Golgi quenched GalT-GFP fluorescence (Movie 1). Subsequently, a substantial fraction of cells exhibited a gradual reappearance of the GalT-GFP signal within newly forming Golgi units (Fig. S4A; Movie 2), reaching a recovery plateau approximately 10–12 hours post-inactivation (Fig. S4B). Notably, the viability of cells undergoing Golgi regeneration appeared largely preserved, as many proceeded through cell division.

### Biogenesis of the new Golgi organelle is associated with changes in its molecular composition

Our data showed that cells could reconstitute new Golgi units after inactivation, but we wondered whether the molecular composition of the recovering Golgi was also restored. We analyzed the localization of a panel of Golgi markers including proteins from various Golgi compartments (*cis*, medial, *trans*) and from different functional classes of Golgi proteins (tethers, coat/adaptors, regulatory GTPases, glycosylation enzymes, structural proteins, etc.). Notably, the panel of markers included proteins with varying topologies ranging from transmembrane to peripheral Golgi proteins.

As previously reported by Jollivet et al. ^9^, GM130 appeared 2 hours post-inactivation at the perinuclear, newly forming Golgi structures flanking the old DAB-positive Golgi (Fig. 2A, empty arrows). Given that GM130 was also detected by EM in early-stage recovering Golgi membranes (Fig. S2A), we used GM130 as a reference marker for the newly forming Golgi to assess the kinetics of other Golgi proteins repopulating the emerging Golgi membranes.

**Figure 2.**
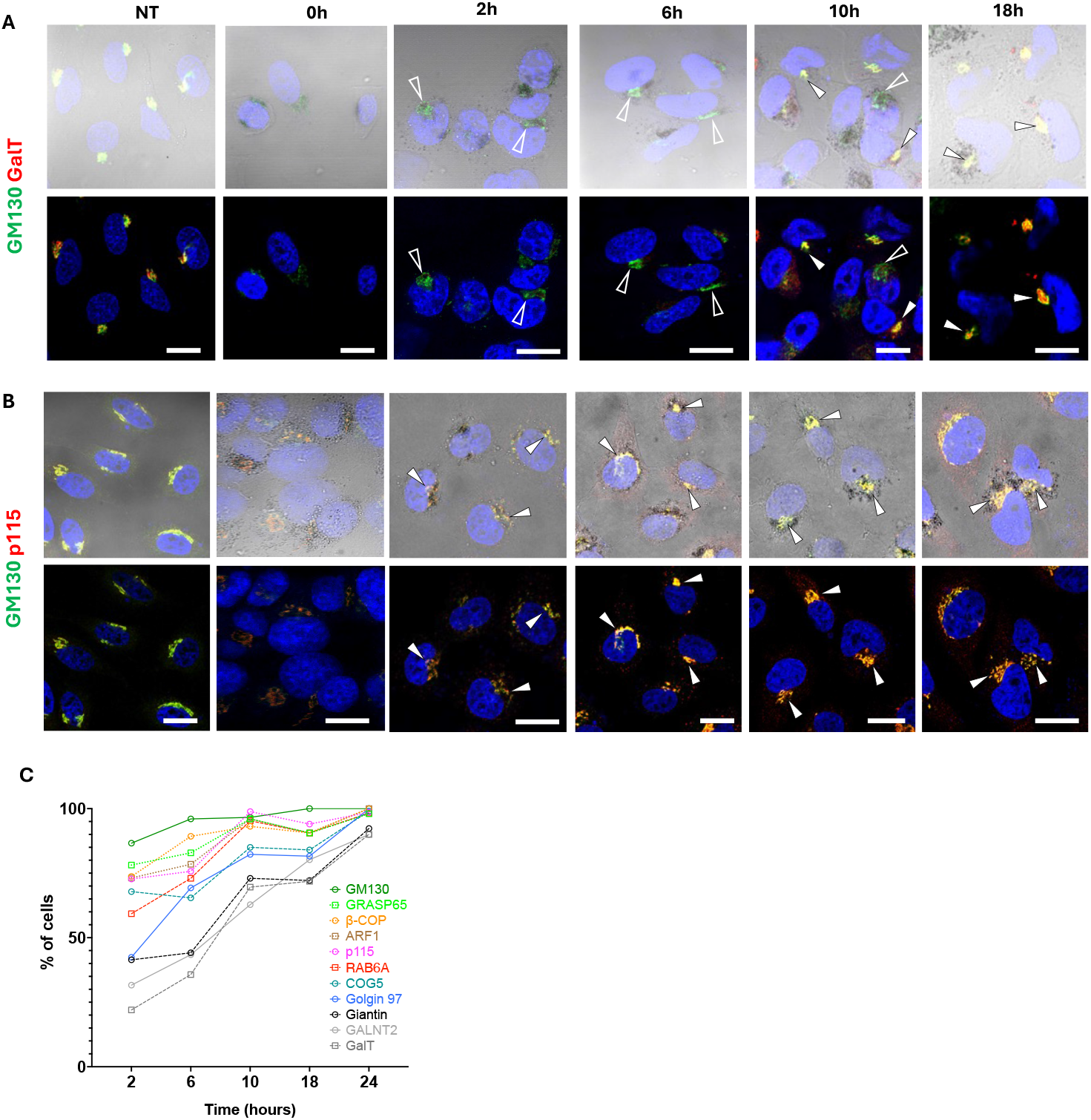
Dynamics of Golgi resident proteins in the newly forming Golgi units. ManII-HRP-expressing HeLa cells were incubated with DAB and H_2_O_2_ to cross-link the preexisting (old) Golgi and fixed for IF immediately or after incubation with fresh medium for 2, 6, 10 or 24 hours. DAB-treated and non-treated (NT) cells were stained for GM130 (green) and with other Golgi marker (red) as indicated in corresponding panels. **A**. Arrowheads indicate newly-forming Golgi which recovered GalT, empty arrowheads show newly forming Golgi without GalT. **B**. Arrowheads show that almost all newly forming Golgi units contain p115. **C**. The graph shows % of cells with corresponding marker in the new Golgi at the different time point after inactivation of the old Golgi. Scale bars: 17 µm (A-C).

We found that Golgi markers appeared in the new Golgi with different kinetics (see examples in Fig. 2A, B and quantification in Fig. 2C). Specifically, transmembrane proteins of the core Golgi region like GalT, Giantin or GALNT2 were detected in a substantial number of cells (>50%) on newly formed Golgi structures only by 10 hours post-inactivation (Fig. 2A, C). Notably, this coincided with the appearance of regular stacks (see EM data in Fig. 1D; Fig. S1D, F), suggesting that a substantial/complete set of Golgi proteins is required to achieve the typical morphology of stacked cisternae. In contrast, the cytosolic proteins p115, GRASP65 and β-COP appeared faster at newly-forming Golgi membranes (Fig. 2B, C) likely due to recruitment from the cytosol and early secretory compartments, such as ERES and the ER-Golgi intermediate compartment (ERGIC), where significant pools of these proteins reside ^14–16^. A similar behavior was observed for ARF1 and RAB6A, likely recruited from the cytosol on Golgi membranes. Notably, even at 24h post-inactivation, a small proportion of the cells did not recover full set of Golgi markers in the newly forming Golgi units (Fig. 2C), indicating that the efficiency of recovery process was heterogeneous across the cell population.

### Newly-formed Golgi supports trafficking of cargo proteins along different routes

We then tested whether newly-forming Golgi structures were functional. We thus combined an analysis of Golgi biogenesis with the evaluation of its trafficking capacities using synchronized RUSH cargoes ^17^.

We first used ManII-HRP cells transfected with a GPI-GFP-RUSH construct, which allowed for monitoring trafficking from the ER along the secretory route to the cell surface. After transfection, cells were incubated with DAB and H_2_O_2_ to inactivate the Golgi apparatus. The cells were then washed and treated with biotin to induce GPI-GFP trafficking. As expected, immediately following the inactivation procedure, GPI-GFP was localized in the ER (Fig. 3A). Two hours later, cells with inactivated Golgi displayed the majority of GPI-GFP within newly formed GM130-positive compartments (Fig. 3A). This indicates that the developing Golgi-like structures were capable of receiving cargo from the ER. This was further confirmed by immuno-EM, which revealed GPI-GFP within vacuolar membranes of newly-forming Golgi (Fig. 3B), which were adjacent to the old DAB-positive Golgi. However, at 2h post-inactivation, we did not observe GPI-GFP at the plasma membrane (PM) of the cells regenerating their Golgi by both light and electron microscopy (Fig. 3A, B). This suggests that the newly forming Golgi units were not fully functional or mature enough to support post-Golgi trafficking. In contrast neighbor DAB-negative cells, whose Golgi remained intact due to low ManII-HRP expression, displayed significant GPI-GFP signal at the cell surface (Fig. 3A, arrowhead). Later, 6h post-inactivation, new Golgi still retained significant amounts of GPI-GFP (Fig. 3A) and only by 10h was able to deliver most of GPI-GFP to the cell surface (Fig. 3A, C) in a substantial number of cells. The activation of effective post-Golgi trafficking of GPI-GFP coincided with appearance of regular stacks of new Golgi (Fig. 3C, D) indicating that functional and structural maturation of the new Golgi correlate in time. Live cell imaging confirmed a significant delay of post-Golgi GPI-GFP trafficking in cells recovering the Golgi after inactivation (compare Movies 3 and 4).

**Figure 3.**
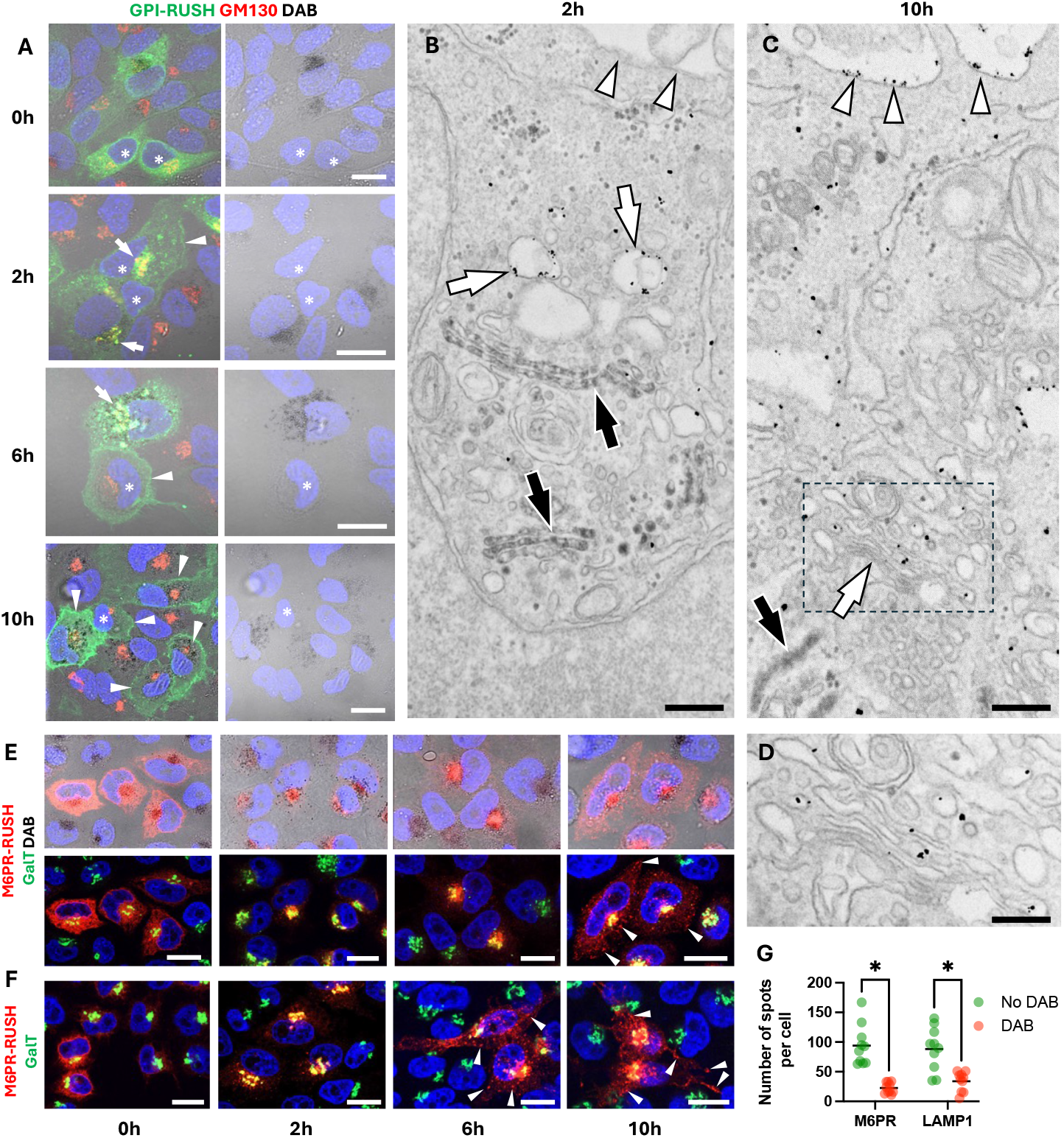
Dynamics of cargo trafficking through newly forming Golgi units. **A**. ManII-HRP-expressing HeLa cells were transfected with GPI-GFP-RUSH cargo reporter and incubated with DAB to cross-link the preexisting (old) Golgi. The cells were fixed immediately or incubated before fixation for 2, 6 or 10h with fresh medium containing biotin to activate GPI-RUSH trafficking and subsequently labelled for GM130. Asterisks indicate cells with no visible DAB, which served as internal control. GPI-RUSH arrived from the ER (0h) to the Golgi of both DAB-positive and DAB-negative cells (2h, arrows). Substantial amount of GPI-RUSH was retained in the newly-forming Golgi of DAB-positive cells (6h, arrows) and was delivered to the plasma membrane (PM) only by 10h postinactivation (10h, arrowheads). No GPI-RUSH retention was detected in the Golgi of DAB-negative cells as GPI-RUSH was efficiently delivered to the PM (2, 6 and 10h, arrowheads) and completely cleaned from the Golgi by 6h postinactivation. **B-D**. Golgi crosslinking and GPI-RUSH trafficking activation was done as described in A. The cells were then prepared for immuno-gold EM to label GPI-GFP-RUSH with anti-GFP antibody. At 2h postactivation GPI-RUSH was detected in the newly-forming Golgi vacuolar membranes (B, white arrows) with no GPI-RUSH in the old Golgi (B, black arrows) or at the plasma membrane (B, arrowheads). At 2h postactivation GPI-RUSH was transported to the plasma membrane (C, arrowheads) and newly formed Golgi stacks were detectable (C, white arrow; see also panel D corresponding to dashed box in C). No GPI-RUSH was detected in the old Golgi (C, black arrows). **E, F**. M6PR-mCherry-RUSH trafficking was activated in ManII-HRP cells with (E) or without (F) Golgi inactivation. M6PR-RUSH was delivered from the ER to the newly forming Golgi but retained there till 6h post-inactivation (E). In contrast, efficient delivery of M6PR-RUSH to post-Golgi compartments (F; 6h, arrowheads) was observed in control cells. In DAB-treated cells cargo reporter appeared in post-Golgi compartments only at 10h after inactivation of the old Golgi (E; 10h, arrowheads). **G**. Quantification of post-Golgi spots containing M6PR-RUSH or LAMP1-RUSH at 6h postinactivation revealed strong reduction in DAB-treated cells, indicating that at this time point newly-forming Golgi was not mature enough to support post-Golgi trafficking. Scale bars: 16µM (A, E, F), 340 nm (B, C), 200 nm (D).

Similar lag in the ability of the new Golgi to export cargoes was observed for proteins directed to endo-lysosomal compartments. We found that in control cells mCherry tagged M6PR or Lamp1 appeared in the post-Golgi endo-lysosomal structures as soon as 2h after release from the ER, while both cargoes were retained within newly forming Golgi units until 10h post-inactivation (Fig. 3E-G). This trend was further confirmed by live cell imaging showing significant delay in M6PR exit from the newly forming Golgi (compare Movies 5 and 6).

Taken together trafficking experiments indicate that developing Golgi requires significant time to acquire functional ability to support trafficking to different post-Golgi destinations. Interestingly, acquisition of this ability correlates in time with appearance of regular Golgi stacks and the recovery of a complete set of Golgi markers.

### De Novo Golgi Biogenesis Requires Transcription

Once we established that cells can rebuild a new functional Golgi, we sought to investigate whether and how they remodel their transcriptome to support *de novo* Golgi biogenesis.

Specifically, we first aimed to determine whether transcription is required for the reconstitution of a novel and functional Golgi complex. To address this, we identified the lowest concentration (1 µg/ml) of actinomycin D (Act-D) that effectively inhibits transcription while maintaining the cell viability for at least 10 hours - the minimum time needed for Golgi recovery (Fig. 4A–C). Act-D treatment resulted in a substantial number of cells in which the newly forming Golgi was hardly detectable after inactivation of the old one (Fig. 4D). In the remaining Act-D-treated cells GM130-positive Golgi units reformed but frequently lacked GalT, indicating structural immaturity (Fig. 4D). This was further mirrored by a marked suppression of GalT-GFP signal recovery in Act-D-treated cells observed in vivo (Fig. S4C, D; Movie 7). At the EM level, Act-D-treated cells failed to regenerate normal Golgi stacks, with most Golgi elements reaching only an incomplete stage of morphogenesis (Fig. 4E-G). These findings suggest that active transcription is essential for the completion of Golgi biogenesis.

**Figure 4.**
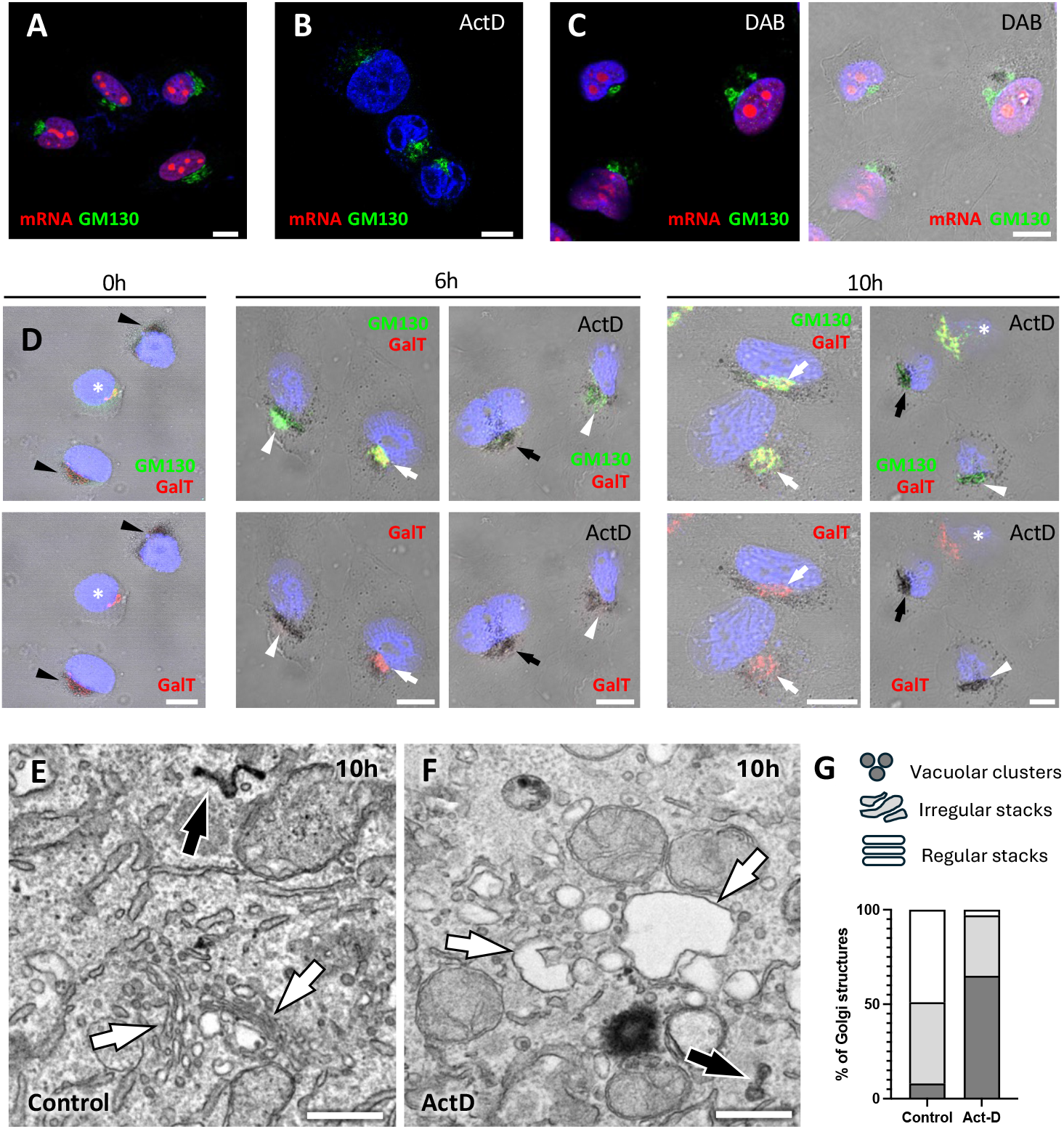
Transcription inhibition affects biogenesis of the new Golgi. A-C. ManII-HRP expressing HeLa cells were labelled to detect newly synthesized mRNA (red), and GM130 (green). Treatment with 1 µg/ml Actinomycin D (ActD) prevented new synthesis of mRNA (B). In contrast, no visible effects on mRNA expression were observed upon treatment with DAB to inactivate Golgi followed by a 6h-recovery period. **D**. Golgi was inactivated by DAB (black arrowheads) and cells were left to recover the Golgi in the presence or absence of ActD. White arrowheads show partially recovered new Golgi containing only GM130. White arrows show that new Golgi containing both GM130 and GalT was recovered only in the cells without ActD treatment. Asterisks indicate the cells, whose Golgi was not inactivated due to the low ManII-HRP expression. Black arrows indicate cells with no detectable GM130-positive new Golgi. **E-F**. The impact of ActD on the architecture of newly forming Golgi was evaluated by EM 10h after inactivation. EM images show that new Golgi (white arrows) did not recover regular stack organization upon ActD treatment (F), as also confirmed by quantification of Golgi phenotypes (G). Scale bars: 8.5µm (A-D), 420 nm (E, F).

This conclusion was further supported by experiments involving cell enucleation, which allowed us to remove the nucleus and generate cytoplasts—nucleus-free cells in which transcription does not occur. While many cytoplasts survived for 8–10 hours (a period sufficient for regeneration of Golgi-like structures in control cells), they failed to rebuild a new Golgi after inactivation of the pre-existing organelle (Fig. S4E, F; Movie 8).

### Single cell RNA-seq reveals transcriptional activation of Golgi genes in cells rebuilding the new Golgi

To determine how transcription contributes to Golgi biogenesis, we investigated changes in the transcriptional landscape of cells during this process. As outlined above, the significant heterogeneity in the kinetics and efficiency of Golgi recovery required the use of single-cell RNA sequencing (scRNA-seq), which allows transcriptome of individual cells to be analyzed during Golgi regeneration process.

ManII-HRP HeLa cells were subjected to Golgi inactivation and then collected for scRNA-seq immediately (0h timepoint) or after incubation with fresh medium for 2, 6, 10, 14, 18 and 24h to allow detailed analysis of transcriptome during different stages of biogenesis of the new Golgi. In parallel, the cells from each time point were subjected to IF and EM analyses to investigate how the dynamics of transcriptional responses correlated with Golgi rebuilding in the framework of the same experiment.

The cells from different time points were prepared for sequencing (see methods) and then pulled together for further NGS analysis. ManII-HRP transcripts were detected in approximately 35000 cells and transcriptomes of individual cells were clustered based on their similarity (Fig. 5A). This analysis revealed that few cell clusters prevailed at early time points post-inactivation (Fig. 5B). Starting from 10h, several cell clusters persisted until 24h, indicating that they might be associated with specific cell states/fates after inactivation of the Golgi apparatus (Fig. 5B). To understand the specific features of these clusters, gene ontology (GO) enrichments analysis for up- and down-regulated genes was performed for each cluster (Fig. 5C; Fig. S5). This analysis revealed that upregulated transcripts in clusters 2 and 10 were enriched in Golgi-associated genes, hereafter referred to as “Golgi genes” (Fig. 5C). Notably, we also found that clusters 2 and 10 shared a significant number of transactivated *Golgi genes* (Fig. 5D). In total, more than 100 up-regulated *Golgi genes* were found in these clusters (Supplementary dataset 1). This number represents a very substantial cohort considering that the sensitivity of scRNA-seq is relatively low as it usually detects about 3000 most abundant transcripts ^18^. Further analysis of individual Golgi genes (*UST, RAB6A* and *COG5*) in UMAP plots indicated that their higher expression was predominantly associated with cellular clusters 2 and 10 (Fig. 5E), thereby corroborating the GO analysis conclusions regarding the upregulation of *Golgi genes* in these clusters.

**Figure 5.**
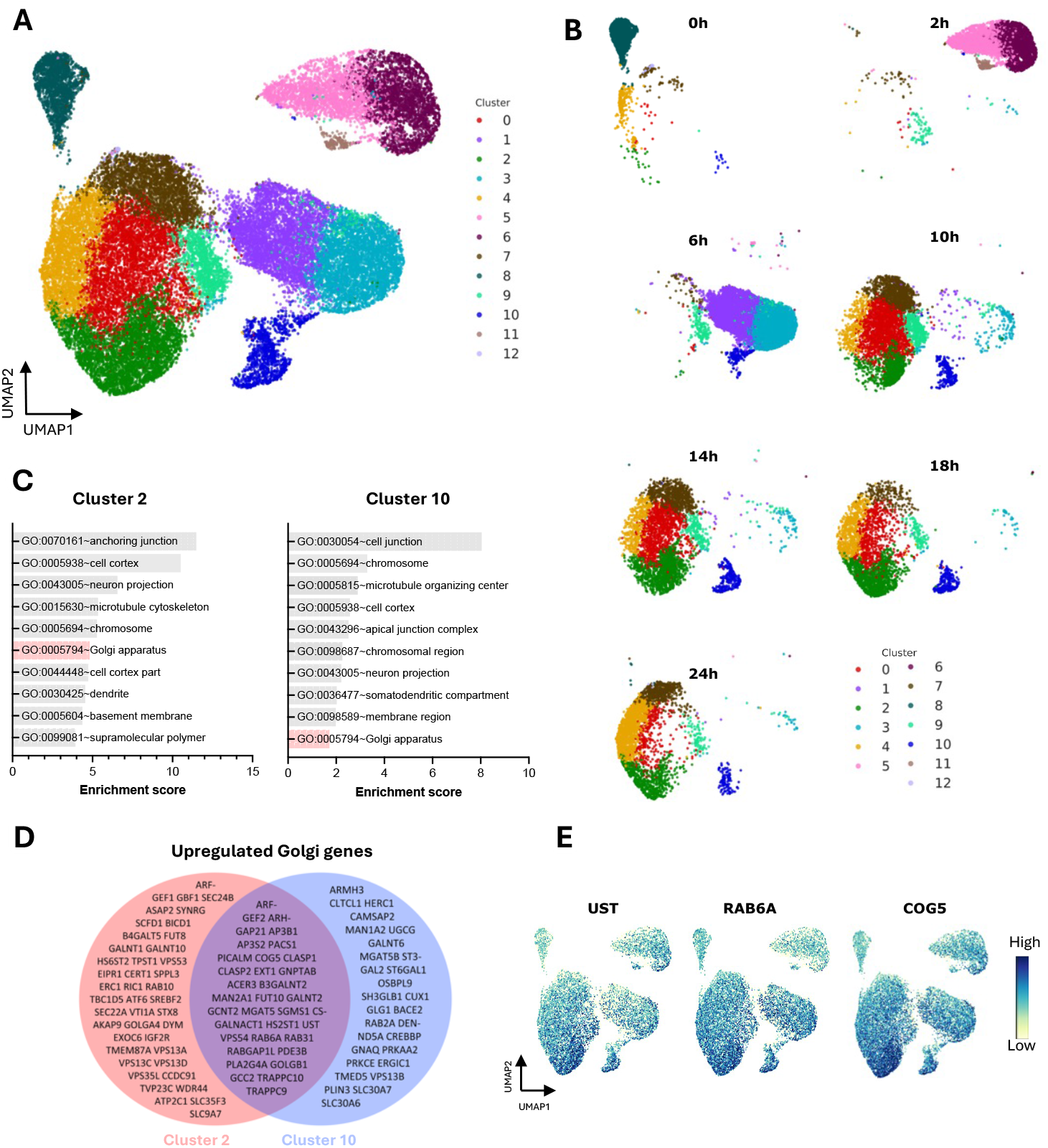
Single cell transcriptomics reveals upregulation of Golgi genes after inactivation of preexisting Golgi organelle. ManII-HRP-expressing HeLa cells were incubated with DAB and H_2_O_2_ to cross-link the preexisting Golgi and collected for scRNAseq immediately or after additional incubation with fresh medium for 2, 6, 10, 14, 18 and 24 h. **A**. UMAP plot shows outcome of the scRNAseq analysis where dots represent individual cells with at least one count for ManII-HRP transcript. The cells were clustered on the basis of similarity of their transcriptomes. **B**. The panel shows UMAP plots for each time point. **C**. Differently expressed genes in each cluster were subjected to gene ontology (GO) enrichment analysis (see also Sup. Fig 8). The graphs show top 10 enriched annotation terms among upregulated transcripts in clusters 2 and 10, including “Golgi complex” (highlighted in red). **D**. VENN diagram shows upregulated Golgi genes that belong to cluster 2, cluster 10, or both. **E**. Expression patterns of Golgi genes *UST, RAB6* and *COG5* are shown in UMAP plots. Cells with higher expression of these genes (dark blue) are concentrated in areas corresponding to clusters 2 and 10 shown in panel A.

Next, we analyzed features of transactivated *Golgi genes* in terms of compartmentalization and function of their protein products. First, we wanted to understand whether specific compartment *cis-*, medial- or *trans*-Golgi prevails among activated genes. Although we found that genes encoding *trans*-Golgi/TGN-associated proteins were more numerous, *cis* and medial *Golgi genes* were also well represented (Fig. 6A), indicating that the activation of the *Golgi genes* in response to Golgi loss is not compartment specific. Second, we analyzed to which functional group do upregulated Golgi genes belong. Figure 6B shows that genes regulating different Golgi functions were represented among activated transcripts. This includes relatively large groups of genes encoding glycosylation enzymes, structural proteins (coats, adaptors and golgins) and proteins regulating membrane trafficking (components of TRAPP, COG, GARP complexes and SNARES). We also found substantial cohort of genes associated with RAB GTPase activities including *RAB6A* (Fig. 6B), whose protein products RAB6A and RAB6A’ regulate numerous Golgi functions ^19^. Other less prominent functional groups of upregulated *Golgi genes* contained genes encoding transporters and cytoskeleton proteins. The group of transporter genes included sugars transporters SLC35A3 and SLC35F3 as well as manganese/zinc transporters SLC30A6, SLC30A7 and ATP2C1 (Fig. 6B), which supply sugar substrates and metal cofactor for glycosylation enzymes residing in the Golgi ^20^. Cytoskeleton-associated genes comprised CLASP1 and CLASP2 (Fig. 6B), whose protein products drive microtubule nucleation at the Golgi ^21^. In summary, this analysis suggests activated Golgi genes were not restricted to a single Golgi compartment or a functional group but targeted more generally Golgi dynamics and function.

**Figure 6.**
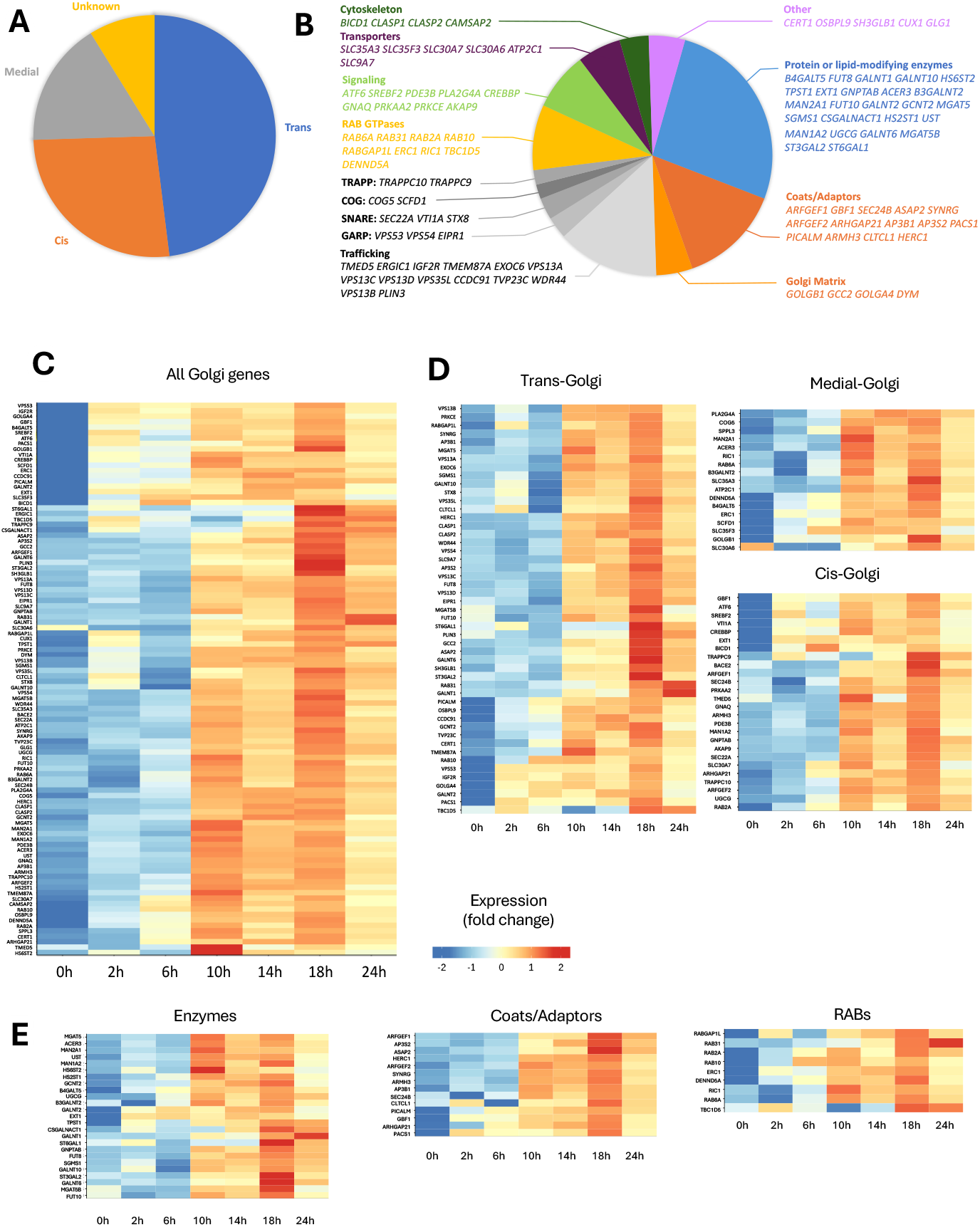
Specific features and expression dynamics of upregulated Golgi genes. **A, B**. Pie plots show proportion of the upregulated Golgi genes, whose protein products are associated with specific Golgi compartment (A) or function (B) **C**. Heat map shows the dynamics of Golgi gene expression after inactivation of the preexisting Golgi organelle. **D, E**. Heat maps indicate that the trend for elevated expression starting from 10h postinactivation is common for majority of the Golgi genes, whose protein products are associated with specific Golgi compartment (D) or function (E).

We also analyzed whether abundance of the protein product somehow impacted whether the corresponding gene is activated after induction of Golgi biogenesis. To this end we mined recently published proteomics data, which provide approximate number of copies per cell for each Golgi protein ^22^. Figure S6A shows that upregulated *Golgi genes* corresponded to proteins ranging from highly to poorly expressed. These comprise *RAB6A*, whose protein product belongs to the top 20 most expressed Golgi proteins, and *GALNT2* encoding a most abundant Golgi glycosylation enzyme ^22^. On the other hand, weakly expressed genes were also transactivated indicating that their protein products might be needed for newly forming Golgi.

### Transactivation of Golgi genes during Golgi biogenesis happens in a synchronous manner

Our scRNA-seq data indicate that extensive Golgi biogenesis after the loss of the preexisting organelle is associated with the activation of a broad set of Golgi-related genes belonging to different functional groups and localized to distinct Golgi compartments. To further characterize this transcriptional response, we examined whether the kinetics of gene activation differ among these groups, such as whether genes associated with the *cis*-Golgi are induced earlier than those related to the *trans*-Golgi, or whether structural components are activated before glycosylation enzymes for example.

Figure 6C illustrates the temporal changes in Golgi gene expression following inactivation of the Golgi apparatus. Notably, most *Golgi genes* showed a significant increase in mRNA levels and in the number of cells expressing these genes, starting between 6 and 10 hours after incubation with DAB (Fig. 6C, Fig. S6B). This elevated expression persisted until 24 hours, with a peak for many genes at 18 hours. This suggests that a broad activation of *Golgi genes* occurs between 6 and 10 hours after the induction of Golgi biogenesis.

Interestingly, a small subset of genes displayed increased transcript levels at earlier time points (as early as 2 hours post-inactivation). This group includes transcriptional regulators such as ATF6, CREBBP, and SREBF2 (Fig. S6C), whose activation may be required to support Golgi biogenesis.

We also investigated whether the activation kinetics of relevant genes differs between Golgi compartments. However, genes associated with the *cis*, medial, and *trans*-Golgi exhibited a similar trend, showing significant activation at 10 hours after the induction of Golgi biogenesis (Fig. 6E). This pattern was also observed across different functional groups of *Golgi genes*, including those encoding glycosylation enzymes, coat/adaptor-related proteins, RAB GTPases and RAB-associated proteins, and signaling factors (Fig. 6F, Fig. S6C).

Importantly, this kinetics of *Golgi gene* activation were not observed in cells lacking detectable ManII-HRP expression (Fig. S6D). This indicates that the Golgi-specific transcriptional response occurred only in cells where the Golgi was inactivated.

### Kinetics of Golgi gene activation correlates with structural maturation of the newly forming Golgi

Our findings suggest that structural and functional transformation of the developing Golgi coincides with the period during which scRNA-seq reveals massive activation of *Golgi genes* (~10 h after inactivation). To further confirm this correlation, we examined Golgi morphology in cells prepared for immunofluorescence and EM, in parallel with cells analyzed by scRNA-seq. New Golgi units in these cells began to acquire a regular stacked morphology by 10 hours after the elimination of the preexisting Golgi, and by 14 hours, substantial number of newly formed Golgi structures were represented by stacks (Fig. S7A-D). Morphogenesis of the Golgi stacks correlated with the appearance of GALNT2 in the emerging Golgi structures (Fig. S7E, F), indicating that these structures recover proteins necessary for their functions, such as glycosylation. This confirms that the activation of *Golgi genes* coincides with the structural and functional maturation of the newly forming Golgi organelle.

We then attempted to directly demonstrate that Golgi gene activation occurs in cells rebuilding the Golgi. This was challenging to monitor using standard bulk methods like real-time PCR, as the cell population following DAB treatment was highly heterogeneous in terms of Golgi recovery efficiency and kinetics. To circumvent this problem, we employed fluorescence *in situ* hybridization (FISH) to visualize mRNA levels of *Golgi genes* in individual cells.

We used ManII-HRP cells expressing GalT-GFP, which allowed us to observe both the DAB-positive old Golgi and the new Golgi marked by GalT-GFP. The cells were subjected to Golgi inactivation and prepared for FISH either immediately or 18 hours after inactivation. The 18-hour time point was chosen because most *Golgi genes* reach peak expression at this time following inactivation of the old organelle (Fig. 6C). To assess Golgi gene expression, RNA-scope FISH probes were selected for genes whose protein products belong to different functional groups, including *UST* and *GALNT2* (glycosylation enzymes), *GOLGB1* (Golgi matrix protein), and *COG5* (membrane tethering protein). These genes were part of the cohort of activated *Golgi genes* identified in the scRNA-seq dataset (Fig. 6B, C).

As shown in Figure 7A, the GalT-GFP signal was quenched by DAB in the Golgi immediately after inactivation, while only a few red fluorescent spots associated with *UST* transcripts were present in the cells. By 18 hours post-inactivation, the fluorescent signal associated with *UST* mRNA had significantly increased in cells recovering the Golgi (Fig. 7A, B). Similar increases in mRNA levels were also observed for *GALNT2, GOLGB1* and *COG5* (Fig. 7A, B), indicating that their transcriptional activation occurs during the biogenesis of the new Golgi. Further control experiments demonstrated that activation of Golgi genes was specifically associated with Golgi biogenesis as incubation with H_2_O_2_ alone fails to stimulate their transcription (Fig. 7C).

**Figure 7.**
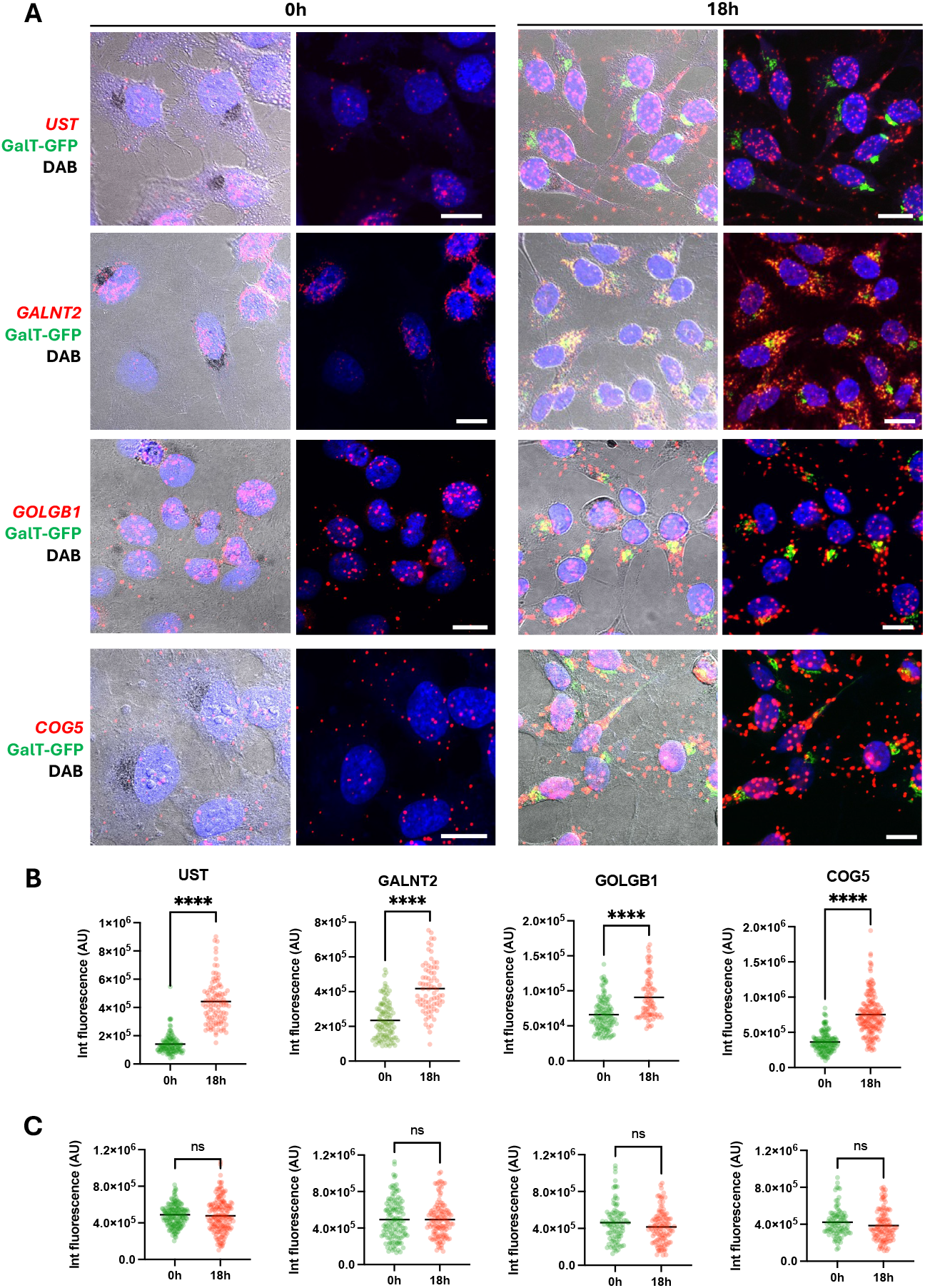
Fluorescence in situ hybridization (FISH) reveals transcriptional activation of Golgi genes in the cells building the new Golgi. HeLa cells expressing ManII-HRP and GalT-GFP were incubated with DAB and H_2_O_2_ to cross-link the preexisting (old) Golgi and fixed immediately or after incubation with fresh medium for 18 hours. The cells were then subjected to FISH with RNAscope probes (see methods) to reveal mRNA of *UST, GALNT2, GOLGB1* and *COG5*. mRNA-associated signal for each gene is represented by red fluorescent spots. Images in panel **A** (0h) show that deposition of DAB polymer in the Golgi quenched GalT-GFP fluorescence, which was recovered in the new Golgi by 18h. The cells recovering the Golgi at 18h show significant increase in red fluorescent signal associated with mRNA of the Golgi genes, as also confirmed by fluorescence quantification in panel **B**. Panel **C** shows no difference in FISH fluorescent signal for the same genes in control cells, which were treated with only H_2_O_2_ and not with DAB. Scale bars: 17 µm (A).

In addition, we examined whether the induction of *Golgi genes* was specific or whether it could be stimulated by the inactivation and subsequent biogenesis of other organelles. To test this, we inactivated endosomes in HeLa cells using HRP uptake (Fig. S8A, B) thereby inducing reformation of the new functional endocytic structures (Fig. S8C). However, endosome biogenesis did not accelerate the expression of *Golgi genes*, which remained unchanged during this process (Suppl. Fig. 8D).

### Induction of Golgi genes during biogenesis requires activity of transcription factor CREB3L1

The massive and concerted induction of *Golgi genes* suggested to us that a biogenesis program was activated under the control of one or a few master genes. We thus hypothesized that key transcription factor(s) (TFs) may be responsible for activating a set of *Golgi genes* and necessary for building the new Golgi.

To identify candidate TFs, we first used the pySCENIC tool ^23^ to infer TF activity from the scRNA-seq data. Then we determined which of predicted TFs might directly regulate Golgi genes, whose upregulated transcripts were enriched in clusters 2 and 10. To this end we used the FIMO tool ^24^ to scan promoter regions of these genes for corresponding TF binding motifs. We additionally included TFs from the CREB3 family in FIMO analysis because (i) these TFs have been reported to regulators secretory pathway-associated genes ^3,25^; and (ii) our set of upregulated Golgi genes included CREBBP, a known transcriptional coactivator of CREB3 proteins ^26^ (see Fig. S6C).

To prioritize predicted TF candidates for further study, we examined their expression profiles in our scRNA-seq dataset (Fig. 8A). TFs expressed in a very limited number of cells, or whose expression declined along the timeline of Golgi recovery, were excluded from further analysis. The remaining candidates (highlighted in pink in Fig. 8A) were subjected to RNAi-based screening. Each TF was silenced using specific siRNAs. The Golgi complex was inactivated, and reassembly of the new Golgi structures was monitored via labeling of GALNT2, which typically marks substantial structural and functional recovery during biogenesis (see Fig. 2C and Fig. S7E, F). We found that silencing SP1, MITF, CUX1, and CREB3L1 significantly inhibited GALNT2 signal recovery in the perinuclear region (Fig. 8B, C). Further validation via EM confirmed severe impairment of Golgi reconstitution in cells silenced for SP1, MITF, and CREB3L1, as mainly vacuolar Golgi precursors can be visualized but not Golgi stacks (Fig. 8D), suggesting that these TFs are involved in *de novo* Golgi biogenesis.

**Figure 8.**
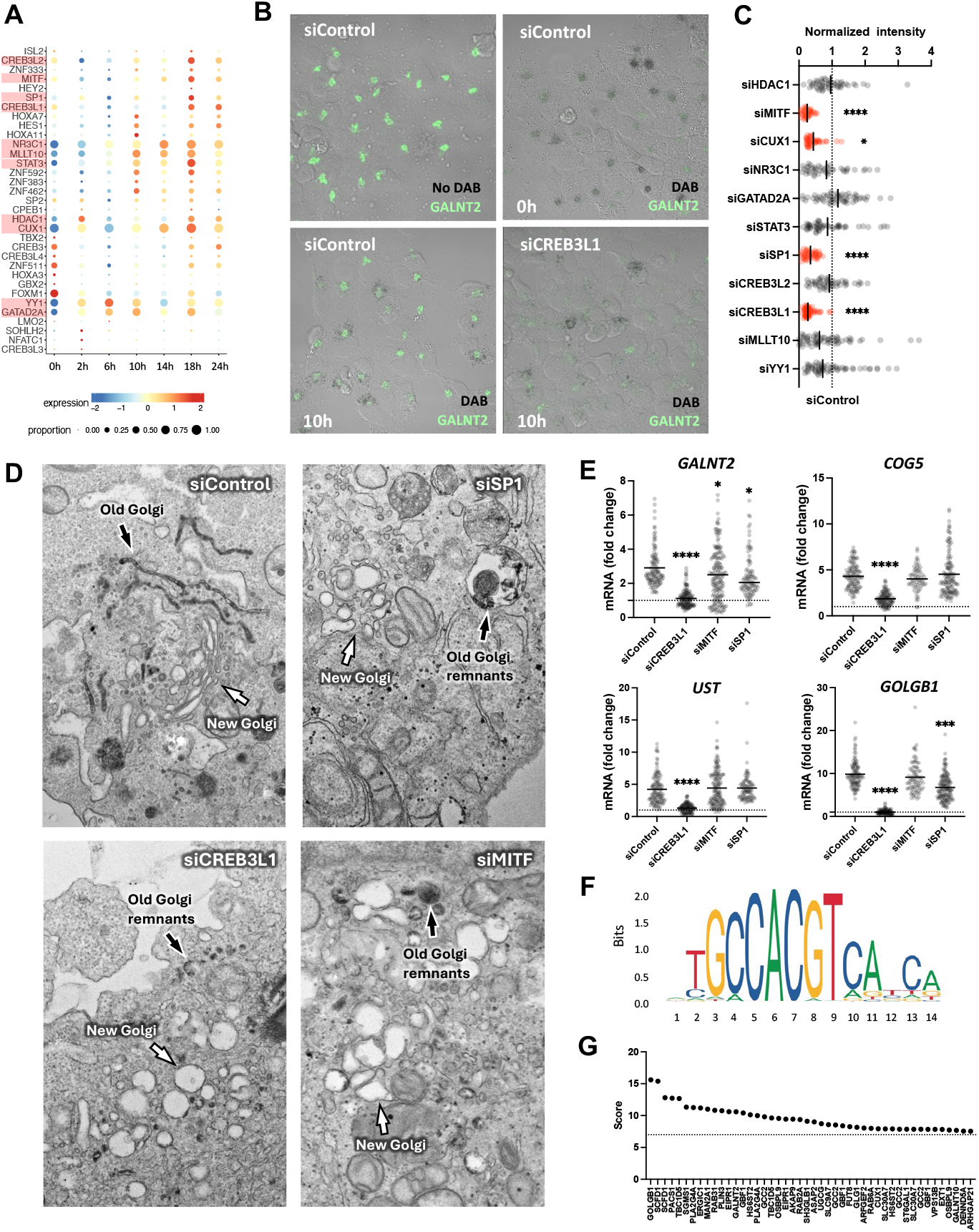
Screening for transcription factors regulating activation of Golgi genes during Golgi biogenesis. **A**. Bubble plot showing the expression patterns of TRANSFAC-predicted transcription factors (TFs) in a scRNA-seq dataset. TF candidates selected for RNAi-based screening are highlighted in pink. **B**. Representative images of control cells treated with scrambled siRNAs (siControl) before inactivation (No DAB), immediately after inactivation (0 h), and 10 h post-inactivation. Efficient GALNT2 recovery is observed in control cells, while CREB3L1-silenced cells show impaired recovery. **C**. Quantification of GALNT2 signal recovery following TF silencing. Values are normalized to siControl (dashed line). siRNAs that significantly reduced GALNT2 recovery are shown in red (^****^p < 0.0001; ^*^p < 0.05; one-way ANOVA; n ≥ 40 cells). **D**. Electron microscopy images showing that control cells reestablish Golgi stacks 10 h after inactivation, whereas cells silenced for SP1, CREB3L1, or MITF fail to do so. White arrows indicate newly formed Golgi structures; black arrows indicate remnants of the old, DAB-positive Golgi. **E**. Expression of GALNT2, COG5, UST, and GOLGB1 was assessed by FISH immediately after Golgi inactivation and again 18 h later. mRNA fold changes between these time points were calculated based on FISH fluorescence intensity and show an increase for all tested genes in control cells (siControl). The graphs depict the impact of transcription factor silencing on this increase for each gene (^****^p < 0.0001; (^***^p < 0.001; ^*^p < 0.05; one-way ANOVA; n ≥ 80 cells). The dashed line indicates gene expression levels immediately after inactivation. **F**. Position weight matrix (PWM) of the CREB3L1 binding motif used for scanning promoter regions of activated Golgi genes. **G**. Motif scores for predicted CREB3L1 binding sites in promoter regions of activated Golgi genes. Higher scores indicate stronger motif matches. The dashed line marks the threshold score of 7, below which binding probability was considered weak. All shown matches were statistically significant (p < 1.0E-04).

Next, we tested whether the inhibitory effects of SP1, MITF, and CREB3L1 silencing on Golgi reconstitution were accompanied by suppression of *Golgi gene* induction. Using FISH in cells recovering Golgi after inactivation, we observed that depletion of CREB3L1 markedly reduced the induction of all tested *Golgi genes* (*UST, GALNT2, COG5*, and *GOLGB1*) (Fig. 8E). In contrast, silencing SP1 or MITF had moderate or no effect on the activation of these genes. These findings suggest that CREB3L1 is a key driver of *Golgi gene* activation during organelle biogenesis, while SP1 and MITF may play more limited or indirect roles.

The Golgi-specific role of CREB3L1 was further confirmed in cells maintained under steady-state conditions (i.e., without Golgi inactivation). Using a panel of *Golgi genes*, we found that CREB3L1 suppression was able to reduce the expression of some of them, while many were unexpectedly induced by MITF and especially SP1 silencing (Suppl. Fig. 9A). In parallel, EM analysis of Golgi ultrastructure revealed significant hypertrophy of pre-Golgi vesicular-tubular membranes at the ERES areas in CREBL1-silenced cells (Fig. 9B) indicating possible impact on the transformation of ER-derived membranes into Golgi cisternae. In contrast no such phenotype was observed in cells exposed to SP1- or MITF-specific siRNAs (Fig. 9B). Finally, we attempted to mimic Golgi gene induction during biogenesis by overexpressing TFs. Figure S9C shows that overexpression of CREB3L1 induced the expression of most tested Golgi genes, whereas SP1 and MITF overexpression induced only a few of them. Together, these data position CREB3L1 as the most compelling TF candidate responsible for *Golgi gene* activation during *de novo* Golgi biogenesis since CREBL1 expression is both necessary and sufficient to induce expression of Golgi genes and biogenesis.

This conclusion was further supported by several lines of evidence. First, bioinformatic analysis of the Golgi gene set identified from the scRNA-seq experiment revealed that promoter regions of a substantial number of these genes contain binding motifs for CREB3L1 (Fig. 8F, G). Second, we found that CREB3L1 was functionally activated during Golgi biogenesis. It has been reported that CREBL1 activation involves proteolytic cleavage, releasing the N-terminal fragment, which translocates to the nucleus and initiates transcription of its target genes ^27,28^. We observed that during Golgi reconstitution, nuclear translocation of CREB3L1 progressively increases and peaks at 6 hours (Fig. S10A, B), preceding the large-scale activation of *Golgi genes*. Finally, bulk RNA-seq of cells overexpressing either full-length CREB3L1 or its constitutively active N-terminal fragment (CREB3L1-CA) showed a significant enrichment of upregulated *Golgi genes* (Fig. S10C, D; Supplementary Datasets 2 and 3), including many that were activated in our Golgi biogenesis experiments (see Supplementary Dataset 1). In contrast, no such enrichment was observed in cells overexpressing MITF or SP1 (Supplementary Datasets 4 and 5), indicating the specificity of CREB3L1 for Golgi genes.

## DISCUSSION

Organelle biogenesis is a fundamental cellular process that enables cells to adapt to environmental cues, developmental signals, and metabolic demands. In this context, Golgi biogenesis plays a central role in supporting increased biosynthetic traffic and expanding Golgi structures during processes such as cell differentiation ^3–6^. While transcriptional regulation of biogenesis has been well characterized for such organelles as lysosomes and mitochondria ^29–31^, the transcriptional control of Golgi biogenesis remains poorly defined.

Most prior studies have examined Golgi-related transcriptional changes in the context of stress responses ^32–35^. However, these models are often confounded by cytotoxicity of chemical stressors, making it difficult to isolate Golgi-specific transcriptional events. An ideal system would enable controlled, synchronous induction of *de novo* Golgi formation, thereby allowing direct interrogation of the associated transcriptional remodeling. Until now, such a model has been lacking.

Although laser ablation has provided mechanistic insight into Golgi morphogenesis ^11,12^, its technical complexity and low throughput preclude large-scale transcriptomic studies. As an alternative, we employed HRP/DAB-mediated inactivation of the preexisting Golgi to trigger synchronized *de novo* organelle biogenesis. While previous use of this technique was insufficient to reconstitute mature Golgi ^9^, we optimized the approach to enable the formation of fully functional Golgi units. Importantly, we demonstrate that this system is well-suited for transcriptome-wide analysis of cells undergoing Golgi reassembly.

### Transcriptional activation of Golgi genes is required for de novo Golgi biogenesis

Using this system, we addressed the previously unanswered question of whether the induction of *Golgi genes* contributes to the morphogenesis of newly forming Golgi organelles. Our finding that massive activation of *Golgi genes* coincides with the structural and functional maturation of nascent Golgi units suggests this may indeed be the case. In line with this notion, we provide direct evidence that transcription is required for Golgi reformation. Inhibition of transcription prevented the formation of mature Golgi stacks, leading instead to irregular, immature Golgi-like structures (see Fig. 4). Similar Golgi precursors can be observed at the early stages of Golgi development, i.e. before massive activation of the Golgi genes (Fig. 1B, Fig. S1B). These precursors contain some Golgi proteins (like GM130, GRASP65, p115, and beta-COP) whose significant pools reside in cytosol and/or ERGIC ^14–16^ from where they can be rapidly recruited to support initial stage of Golgi morphogenesis. However, in the absence of active transcription, these nascent structures failed to mature into functionally competent and normally shaped, Golgi complexes.

This suggests that later stages of biogenesis—particularly those restoring Golgi stacking, trafficking, and enzymatic function—require robust transcriptional input. Indeed, we observed a strong correlation between transcriptional activation of Golgi genes at ~10 hours post-inactivation and the emergence of functional, stacked Golgi structures. This timeline closely mirrors Golgi recovery kinetics following laser ablation ^11,12^, reinforcing the notion that loss of Golgi triggers a transcriptional program essential for its reconstruction.

Another key question we addressed was whether cells require a broad activation of Golgi genes to rebuild the organelle, or only the activation of a compartment- or function-specific subset, to rebuild the organelle. We also asked whether activation of such subsets occurs in a sequential manner.

Previous studies suggest that Golgi assembly after chemical disruption proceeds in a cis-to-trans order ^36,38^. Therefore, we did not rule out the possibility that the cell might first activate cis-Golgi genes, followed by genes associated with the trans-Golgi compartment.

However, we observed that the activated genes spanned diverse Golgi compartments and functional classes, suggesting that Golgi regeneration is orchestrated as a coordinated transcriptional program rather than a stepwise activation of specific modules. Genes encoding cis- and trans-Golgi proteins, as well as those involved in glycosylation, trafficking, and ion transport, were all upregulated with similar kinetics.

This broad activation supports the idea of a unified Golgi regulon, enabling the simultaneous expression of components required for the organelle structural and functional integrity. Such coordination is likely critical for processes like glycosylation, which requires the synchronized activity of enzymes localized to distinct sub-Golgi regions, along with transporters supplying substrates and ion cofactor ^20,37^. In addition, these components may be critical for cisternal morphogenesis, as the absence of their expression prevents the transformation of vacuolar membranes in Golgi precursors into the flattened disks characteristic of mature cisternae. Thus, our findings suggest that Golgi biogenesis is driven by a highly integrated transcriptional mechanism, fine-tuned to ensure that the growing organelle acquires normal architecture and full functional capacity.

### Is CREB3L1 the Master Regulator of the Golgi Gene Network?

Organelle-specific gene networks are typically governed by master transcriptional regulators—for instance, TFEB and TFE3 for lysosomes ^29,30^, or PGC1α for mitochondria ^31^. The coordinated activation of Golgi-associated genes during *de novo* Golgi biogenesis suggests the existence of a similar regulatory mechanism for the Golgi apparatus.

We identified three transcription factors—SP1, MITF, and CREB3L1—whose silencing significantly impaired Golgi reassembly. CREB3L1 was particularly relevant as knockdown robustly suppressed the induction of all tested Golgi genes in cells undergoing Golgi reconstruction (see Fig. 8). In contrast, SP1 and MITF silencing inhibited only some of these genes and with markedly lower magnitude. This points to a more selective and central role for CREB3L1 in Golgi gene activation during organelle biogenesis.

Golgi specificity was further supported by CREB3L1 gain- and loss-of-function experiments. Overexpression of CREB3L1 (or its constitutively active variant) induced *Golgi gene* expression, while its knockdown led to repression of some *Golgi genes*. Notably, SP1 or MITF did not produce comparable effects, as their silencing in some cases paradoxically increased expression of *Golgi genes*. These findings suggest that while SP1 and MITF may contribute to Golgi biogenesis, their roles are likely indirect or involve additional, yet undefined, mechanisms.

CREB3L1 is a member of the CREB3 transcription factor family, several of which have been implicated in transcriptional regulation of secretory pathway genes, including those involved in ER-to-Golgi trafficking and Golgi function ^3,28,39^. Previous studies have shown that CREB3L1 and its homolog CREB3L2 are activated in response to increased secretory load, and are required for the transcriptional upregulation of genes supporting Golgi expansion ^3,40,41^, further supporting our findings.

Promoter analysis revealed putative CREB3L1-binding sites in a substantial number of *Golgi genes* upregulated during biogenesis (Fig. 8F, G; Suppl. Table 1). Interestingly, some CREB3L1-responsive genes (e.g., *COG5, UST*) lacked predicted CREB3L1 motifs, yet their activation was still inhibited by CREB3L1 silencing. This suggests the possibility of indirect regulation or the involvement of non-canonical binding sites not yet annotated in current databases.

Taken together, our data position CREB3L1 as a key transcriptional driver and candidate master regulator of Golgi biogenesis, able to coordinately activate a broad set of Golgi-associated genes during organelle formation.

### How does CREB3L1 sense the cellular need to activate Golgi genes?

We found that the biogenesis of a new Golgi apparatus coincides with increased nuclear translocation of CREB3L1 (Fig. S10A, B). Such translocation is a known marker of ER stress, even in its mild form ^27^. ER stress stimulates CREB3L1 cleavage by Site-1 and Site-2 proteases leading to consequent releases of the N-terminal fragment, which translocates into the nucleus and activates the transcription of its target genes ^27,28^.

We propose that ER stress during new Golgi formation may be triggered by a transient “traffic jam” in the early secretory pathway, caused by immature Golgi precursors failing to efficiently sort and export cargo. In this context, the limited capacity of the nascent Golgi to retrieve ER-resident chaperones may be sufficient to activate the unfolded protein response, leading to CREB3L1 activation. The transcriptional activity of CREB3L1 towards *Golgi genes* would then support the functional maturation of the new Golgi thereby enhancing the capacity of the secretory system. In line with this hypothesis, CREB3L1 activation precedes the structural and functional maturation of the Golgi. Nuclear translocation of CREB3L1 begins 2 hours post-activation and peaks at 6 hours, anticipating the massive induction of target Golgi genes observed at 10 hours—when the Golgi begins to acquire functional and structural maturity.

### Transcriptional program of Golgi biogenesis mirrors physiological secretory remodeling

It is important to note that, in addition to CREB3L1/2, several TFs have been proposed to regulate Golgi-associated genes. These factors typically emerge from studies analyzing transcriptional responses under so-called Golgi stress, induced by chemical agents that disrupt Golgi architecture and function. However, the identity of the activated Golgi gene sets and their regulatory TFs varies significantly across studies, suggesting that Golgi stress responses are highly context-dependent and shaped by the nature of the stressor.

In this study, we sought to determine whether mechanisms implicated in the Golgi stress response also contribute to Golgi biogenesis. When comparing gene sets activated by three distinct Golgi stressors—Brefeldin A (BFA), Monensin, and Golgicide ^35^—to the set of genes activated during Golgi biogenesis in our model, we observed minimal overlap (Fig. S11A). A similarly low degree of overlap (Fig. S11B) was observed with genes activated in response to the Golgi stressor OSW-1 ^42^. These findings indicate that the transcriptional programs underlying Golgi biogenesis and Golgi stress responses are largely distinct. Similar to the ER, the Golgi harbors multiple stress-responsive pathways ^33^. Indeed, Golgi stress has been shown to activate TFs such as MLX, TFE3, ELK1, and CREB3 ^32–35^. Among these, only CREB3 belongs to the same family as CREB3L1, which we identified as a specific regulator of the Golgi gene network during biogenesis. However, whether CREB3 activates the same network is unknown, as information about its Golgi gene targets under BFA-induced stress conditions remains limited to *ARF4* ^34^.

Interestingly, the transcriptional program activated during *de novo* Golgi biogenesis appears to resemble the gene expression changes observed during cell differentiation, where there is a well-documented need for expanded Golgi capacity to support increased secretory activity. For instance, during differentiation of naïve B cells into antibody-secreting plasma cells, the Golgi drastically expands nearly sevenfold ^4^, accompanied by a robust induction of Golgi-associated genes ^43,44^. Remarkably, nearly half of the Golgi genes we identified during Golgi biogenesis overlapped with those induced during B cell differentiation (Fig. S11C). A significant overlap was also observed with Golgi genes upregulated during differentiation of endometrial stromal cells (Fig. S11D), which remodel the Golgi to support secretion of implantation-related factors ^40^. Notably, gene activation in both differentiating B cells and endometrial cells requires the activity of CREB3L1 and/or CREB3L2 ^40,41^.

These findings suggest that the transcriptional program governing Golgi biogenesis during organelle reassembly is shared with physiological states requiring secretory adaptation. This is further supported by reports implicating CREB3L1 in Golgi expansion during differentiation of thyroid epithelial cells and neurons ^3,26^. Collectively, our results argue that Golgi biogenesis is regulated by a specific transcriptional program that operates not only during organelle regeneration but also in biologically relevant contexts of cellular differentiation and response to stimuli activating secretion.

## ACKNOWLEDGMENTS

We would like to acknowledge C. Alvarez for CREB3L1 constructs; M.A. De Matteis and C. Settembre for critical reading of the manuscript; Lorenzo Vaccaro for help with preparation of lentiviruses; TIGEM core facilities (Histopathology, Bioinformatics, Microscopy and Imaging) for technical and analytical support.

## Authors contributions

Conceptualization – RSP, AK, JDB, FP; Data collection – FF, DA, EVP, FR, RC, JS, RP, MS, NCS, JV; Data analysis - FF, DA, EVP, FR, RP, XBC, RDC; Data curation - XBC, RDC; Writing original draft – RSP, FF, DA; Writing, draft editing – FP, BG, AK, XBC, RDC, NCS; Funding acquisition – RSP, AK, JDB, FF, FP.

## Data and materials availability

ScRNA-seq data and bulk RNA-seq data were deposited to GeneExpression Omnibus (GEO) under accession codes GSE288395 and GSE299965, respectively. All other data are available in the manuscript or the supplementary materials.

