## Supplementary text and figures for "*De novo* Golgi biogenesis requires coordinated transactivation of a Golgi regulon"

**The PDF file includes:**

Materials and Methods
Supplementary Text
Figs. S1 to S11
References

### MATERIALS AND METHODS

#### Antibodies, plasmids and other reagents

Primary antibodies against flowing proteins were used in the study: BetaCop (Abcam, ab2899), COG5 (Abnova, PAB21288), TGN46 (Bio-Rad, AHP500), GM130 (BD Biosciences, 610823), GALNT2 (Sinobiological, 13764-T62), Syntaxin5 (Synaptic System, 110 053), ARF1 (Proteintech, 10790-1-AP), p115 (Proteintech, 13509-1-AP), RAB6A (Proteintech, 10187-2-AP), GRASP65 (Thermo Fisher Scientific, PA3-910), Giantin (kindly provided by Dr. M.A. De Matteis, TIGEM, Pozzuoli, Italy), GFP (Abcam, ab290). Secondary anti-rabbit, anti-mouse, and anti-sheep antibodies conjugated with Alexa Fluor 488 or 568 were purchased from Life Technologies (Carlsbad, CA). GPI-GFP-RUSH (Str-KDEL_SBP-EGFP-GPI) DNA constructs for trafficking experiments was reported previously (*1*). LAMP1-mCherry-RUSH (Str-KDEL_IRES_SBP-mCherry-Lamp1) and M6PR-mCherry-RUSH (Str-KDEL_IRES_SBP-mCherry-CD-M6PR) constructs (*2*) were obtained from Addgene. CREB3L1 construct was kindly provided by Dr. Cecilia Alvarez (Universidad Nacional de Córdoba, Córdoba, Argentina). RNA Synthesis assay kit was obtained from Abcam (ab228561). pEGFP-N1-MITF-M (Addgene # 38131) and CMV_SP1_HA (VectorBuilder Inc, Vector ID VB250121-1102aju) were used to overexpress MITF and SP1 in HeLa cells.

#### Cell culture and cytoplast generation

HeLa ManII-HRP cells, were cultured in Dulbecco's Modified Eagle Medium (DMEM) supplemented with 10% fetal bovine serum (FBS), penicillin/streptomycin, and 2 mM L-glutamine. HeLa ManII-HRP cell line stably expressing GalT-GFP were generated using transduction with lentiviral particles. pLVX-EF1a-GalT-IRES-Puromycin cDNA plasmid (Addgene #134862), was used to generate lentiviral particles using a second-generation packaging system in X293T (Takara Bio) cells. HeLa ManII-HRP cells were transduced with lentiviral particles at a multiplicity of infection (MOI) 30, in the presence of 8 μg/mL polybrene (Sigma-Aldrich) to enhance transduction efficiency. Cells were incubated with the virus-containing medium for 24 hours, after which the medium was replaced with fresh culture medium. 48–72 hours post-infection, transduced cells were selected with puromycin.

Generation of nuclear-free cells (cytoplast) was performed by centrifugation of cells grown on glass coverslips (Prescott et al., 1972). Thirty minutes prior to centrifugation, cytochalasin D (Sigma-Aldrich, C8273) was added to the culture media at 5-mg/ml final concentration. Coverslips were centrifuged at 10,000 RPM in a FiberLite F21-8X50 rotor (Sorvall) for 30 minutes at 35°C in media containing cytochalasin D. Immediately after centrifugation, the coverslips were rinsed four times with culture media and incubated in fresh DMEM/F12 GlutaMax medium supplemented with 10% FBS for 1 h at 37°C in 5% $CO_2$ prior to DAB-mediated inactivation of the Golgi.

#### RNA Interference

Gene silencing was performed using siRNAs delivered with Lipofectamine™ RNAiMAX Transfection Reagent (Thermo Fisher Scientific, Cat. No. 13778075), according to the manufacturer's instructions. Cells were transfected with siRNA complexes prepared in Opti-MEM™ Reduced Serum Medium. For each target gene, transfections were carried out using a combination of two or three distinct siRNA sequences to maximize silencing efficiency and minimize off-target effects. Final siRNA concentrations and transfection conditions were optimized based on preliminary titration experiments. Cells were processed for downstream analyses 48–72 hours post-transfection, depending on the assay. Silencing efficiency was assessed by qRT-PCR. All siRNA sequences used in this study are listed in Supplementary Dataset 2.

**DAB-mediated Golgi inactivation**

The 10× DAB stock solution (D8001, Sigma-Aldrich) was prepared by dissolving 50 mg of 3,3'-diaminobenzidine (DAB) in 0.1 M HCl. The working solution (1× DAB) was obtained by diluting the stock solution in 0.1 M Tris-HCl (pH 7.4). Cells expressing ManII-HRP were incubated with 1× DAB and 0.03% $H_2O_2$ at room temperature at the light microscope stage. The appearance of brown DAB signal within the cells was tightly monitored by bright field microscopy to avoid DAB deposition outside the Golgi. Once the presence of the light-brown signal in the Golgi was evident (usually after 3-5 min) the cells were washed with phosphate-buffered saline (PBS) and transferred to pre-warmed complete medium (37°C) to stimulate formation of the new Golgi organelle and to analyze this process with different approaches (see below).

**Immunofluorescence (IF)**

To assess the morphology and composition of the newly-forming Golgi the cells were fixed at different time points after DAB-mediated inactivation with 4% paraformaldehyde (PFA) in PBS for 10 minutes, followed by a 30-minute incubation in a blocking/permeabilization solution containing 1%BSA, 0.5% saponin and 50 mM $NH_4Cl$ in PBS. Primary antibodies were diluted in the blocking/permeabilization solution and incubated with cells for 1 hour or overnight. Secondary antibodies were similarly diluted and incubated for 45-60 minutes. The labelled cells were mounted in Mowiol (Sigma, D2522) and examined using a ZEISS LSM 700 or LSM 800 confocal microscope using a 63× 1.4 NA oil immersion objective and appropriate settings of excitation and emission for image capture. For each Golgi marker the percentage of cells with marker-positive new Golgi was calculated at each time point after DAB-mediated inactivation.

**Electron microscopy (EM)**

Ultrastructure of the newly-forming Golgi was analyzed in ManII-HRP HeLa cells at different time points after DAB-mediated inactivation. For immuno-electron microscopy analysis, the cells were prepared as described previously (*3*). Briefly, after fixation with mixture of 4%PFA and 0.05% glutaraldehyde in 0.2 M HEPES for 15 min and with 4%PFA alone for 30 min, the cells were incubated with blocking/permeabilizing solution: 0.5% bovine serum albumin (BSA), 0.1% saponin, 50 mM $NH_4Cl$ in PBS for 20-30 min. Primary antibody against either GM130 (BD Biosciences, Franklin Lakes, NJ, USA) or GFP (Abcam, Cambridge, UK) were diluted in blocking/permeabilizing solution and added to the cells overnight, followed by 2h incubation with 1.4nm gold-conjugated Fab' fragment of anti-rabbit IgGs (Nanoprobes, Yaphank, NY, USA). GoldEnhance™ EM kit (Nanoprobes, Yaphank, NY, USA) was used to enhance ultrasmall gold particles. Then cells were scraped, pelleted, post-fixed in OsO4 and uranyl acetate and embedded in Epon.

For routine EM, the samples were fixed in 1% glutaraldehyde in 0.2 M HEPES for 60 min and post-fixed and embedded as indicated above. From each Epon-embedded sample, thin 60-nm sections were cut using a Leica EM UC7 (Leica Microsystems, Wetzlar, Germany). EM images were acquired from thin sections under Tecnai G2 Spirit BioTwin electron microscope (Thermofisher, Eindhoven, The Netherlands) equipped with a VELETTA CCD digital camera (Soft Imaging Systems GmbH, Munster, Germany). Inactivated Golgi compartments and their remnants were identified based on the presence of electron-dense DAB polymer. Newly-forming Golgi units were identified in immune-EM specimens based on (i) the presence of GM130-associated gold particles and (ii) absence of DAB. The structures with similar ultrastructural features in routine EM specimens were considered as newly-forming Golgi membranes and were arbitrary divided into three main classes vacuolar clusters, irregular stacks and regular stacks for morphometric analysis.

**RUSH assay of membrane trafficking**

Ability of the newly-forming Golgi to support trafficking of different cargo proteins was evaluated with RUSH (Retention Using Selective Hooks) approach. LipoD293™ (SignaGen Laboratories, Frederick, MD, USA) or Lipofectamine 3000 (Thermo Fisher Scientific) were used to transfect RUSH cDNA constructs encoding reporters directed to either plasma membrane (GPI-GFP-RUSH) or endo-lysosomal compartments (M6PR-mCherry-RUSH, LAMP1-mCherry-RUSH). RUSH cargoes release from endoplasmic reticulum in control or DAB-inactivated ManII-HRP HeLa cells was induced by adding pre-warmed Leibovitz medium (Life Technologies) supplemented with 40 µM biotin (Sigma-Aldrich, Cat. No. B4501). The efficiency of RUSH cargo trafficking to different compartments of the biosynthetic route was assessed by IF or immuno-EM in cells fixed at the different intervals after release of the reporters from the ER. The analysis of M6PR-mCherry-RUSH or LAMP1-mCherry-RUSH delivery to target destination was conducted based on quantification of number of RUSH cargo-positive post-Golgi structures per cell.

**Live-cell microscopy**

Cells were grown on #1.5 glass coverslips in Petri dishes for 48-72 h. After DAB-mediated inactivation of Golgi, phenol-red free mixture of DMEM/F-12 containing 10% FBS was added to the cells, and coverslips were mounted on Rose chambers and placed on the microscope stage. The chambers were maintained at 37.0 ± 0.3°C within a custom-built enclosure. Golgi regrowth and trafficking were acquired in multi-mode (differential interference contrast [DIC], bright-field, and wide-field fluorescence) time-lapse recordings on a Nikon TE2000-E2 microscope equipped with a PlanApo 60X NA 1.4 oil-immersion DIC H objective (Nikon) and LED illuminator (CoolLED *p*E-4000). Recordings were done at 10-15 min intervals at 1 mm z-steps and captured on an Andor Zyla (VSC-04182) camera at 0.22 mm XY pixel size. The system was controlled by NIS-Elements Software.

All time-lapse recordings acquired with Nikon TE2000-E2 microscope and used for quantification, were analyzed using Fiji. For the quantification of GalT-GFP, all time-lapse recordings were acquired using identical illumination settings. The mean pixel values of GalT-GFP were calculated from maximum-intensity projections (MIP) of z-stacks with a region-of-interest (ROI) drawn around the Golgi at each timepoint. The Golgi areas were identified based on DAB deposition observed in the bright-field channel at each timepoint. The mean pixel values were then corrected by subtracting out the background signal.

**Endocytic HRP uptake and endosome inactivation**

HeLa cells were incubated for 90 min at 37°C in DHB (DMEM + 20 mM HEPES + 0,1% BSA). Then 2mg/mL HRP in DHB was added for 30 min at 37°C, whereupon cells were washed 4 times with cold PBS to eliminate the Extracellular HRP. After HRP uptake the cells were fixed to assess HRP colocalization with endocytic markers (EEA1 or LAMP1) by confocal microscopy or subjected to inactivation of HRP-loaded endosomes. For inactivation the cells were incubated in DAB solution (0,1 M Tris HCL pH 7,4 + 0,25 mg/mL DAB + 0,02% H2O2) for 2 min at room temperature, then were washed in PBS and fixed or re-incubated with fresh medium. To check recovery of functional endosomes, at 18h post-inactivation HRP was re-added to the cells for 30 min. The cells were then fixed and the efficiency of HRP uptake was assesses by its IF labeling.

**RNA synthesis assay**

Efficiency of RNA-synthesis was evaluated with RNA Synthesis assay kit (Abcam). ManII-HRP HeLa cells were incubated with RNA Label component of the kit for 1h at 37°C. Then the cells were fixed and permeabilized according to the manufacture protocol. RNA Reaction cocktail was added to the cells for 30 minutes at room temperature in dark. The cells were then washed three times with Wash Buffer and stained with DAPI for nuclei and antibody of interest (when needed). Red fluorescent signal associated with de novo synthesized RNA was detected using excitation and emission settings for AlexaFluor568 at

ZEISS LSM800 confocal microscope. Transcription inhibitor Actinomycin-D was used as a control to evaluate the specificity of the labelling.

#### Fluorescent In Situ Hybridization (FISH)

To analyze mRNA levels of Golgi-associated genes, we customized the RNAscope™ FISH protocol (Advanced Cell Diagnostics, USA) based on transcript size, following the manufacturer's guidelines. RNA in situ hybridization was performed using the RNAscope Multiplex Fluorescent v2 reagent kit (ACD, no. 323285). After Golgi inactivation formalin-fixed ManII-HRP cells were pre-treated with RNAscope Protease III (ACD, no. 322380), diluted in PBS at room temperature, according to a customized protocol designed to preserve the endogenous GFP signal. After pre-treatment, each well was incubated with different probes for corresponding gene (*COG5* #1297181-C3, *GALNT2* #1053611-C3, *GOLGB1* #1297191-C2, *UST* #851011-C3), followed by the amplification cascade as outlined in the manufacturer's protocol. Detection was performed using TSA Vivid Fluorophore 570 (ACD, no. 323272). Finally, cells were counterstained with DAPI, mounted with GEL/MOUNT (Biomeda, Foster City, CA, USA), and analyzed using a ZEISS LSM900 confocal microscope. Intensity of the FISH labelling per cell was quantified using Fiji software.

#### RT-qPCR

To evaluate gene expression levels in silencing and overexpression experiments total RNA was isolated using the RNEasy Mini Kit (Qiagen, Germantown, MD, USA), from treated and control cells. After that, reverse transcription of RNAs was performed using primers indicated in Supplementary Dataset 3 and QuantiTect Reverse Transcription kits (Qiagen). RT-qPCR experiments were performed using Light Cycler 480 Syber Green I Master Mix (Roche, Basel, Switzerland) using a LightCycler 480 II Real-Time System (Roche) in 96-well plates. RT-PCR results were analyzed using the $2^{-\Delta\Delta Ct}$ method, normalized against the housekeeping *ACTB* gene encoding β-actin.

#### Single cell RNA sequencing (scRNA-seq)

***Sample collection*.** Cells were subjected to DAB-mediated Golgi inactivation as described above. After removal of the DAB solution, the samples were washed in PBS and collected for RNA-seq immediately, or after different incubation intervals (2, 6, 10, 14, 18 or 24 hours). For each time point approximately 100000 cells were collected and their viability was checked by trypan blue staining to ensure that the percentage of dead cells does not exceed 5%. In parallel from each time point the cells were prepared for EM and IF analysis to assess the degree of morphogenesis of the newly-forming Golgi.

***Sequencing*.** Libraries were prepared by using the NEGEDIA NGD0050R_single-cell RNA-seq (10X Genomics) Sequencing service (Negedia s.r.l). In brief, single-cells were suspended in phosphate-buffered saline containing 0.04% bovine serum albumin, filtered using 40 um cell strainer (Biologix), and their concentration was evaluated at LUNA-II™ Automated Cell Counter (Logos Biosystems). The cell suspension was loaded onto the Chromium Single Cell G Chip Kit (10x Genomics) and run on the Chromium Single Cell Controller (10x Genomics) to generate single-cell gel beads emulsion, according to the manufacturer's protocol. The single-cell 3' Library and Gel Bead Kit V3.1 (10x Genomics) was used to generate cDNA and the final libraries. The cDNA quality was assessed using High sensitivity D5000 screen tape on the Agilent 4200 TapeStation system (Agilent Technologies). The quality of libraries was assessed by using screen tape High sensitivity DNA D1000 (Agilent Technologies). Finally, the libraries were sequenced on a Novaseq6000 sequencer (Illumina) according to the manufacturers' specifications.

***Data pre-processing*.** Demultiplexing was performed with Cell Ranger (*4*) (v. 7.1.0), with default settings. Then, a customized reference genome was generated to identify ManII-HRP positive cells by merging the human GRCh38 genome from Ensembl and the ManII-HRP sequence. A customized .gtf file was also generated. Mapping and counting were performed using the Cell Ranger (*4*) counting function (v. 7.1.0) with predefined parameters and using the customized genome as a reference. Then, 3 files (matrix.mtx.gz, features.tsv.gz and barcodes.tsv.gz) were generated for each sample, which formed the input for the Seurat pipeline (*4*).

***Data analysis.*** Data were analyzed in Rstudio (R version 4.3.0) using the Seurat package [2] (version 4.4). A Seurat object, including only ManII-HRP positive cells was generated (cells containing at least 1 count for ManII-HRP). Cells that expressed more than 500 genes, had more than 1000 counts and less than 15% of mitochondrial gene expression were filtered in. A post-filtered dataset of 13092 genes expressed in 35866 cells was used for downstream processing. Count data was then processed using SCTransform (*5*), which normalizes and stabilizes variance through a regularized negative binomial regression and regressed out against the percentage of mitochondrial genes. Using FindVariableFeatures, the 2000 most variable features were identified and then the Principal Component Analysis (PCA) was performed on these features using the runPCA function. Cell clusters were determined via Shared Nearest Neighbor (SNN) and k-Nearest Neighbor (KNN) analyses using the FindNeighbors function with the first 6 Principal Components and FindClusters function with resolution 0.5 respectively. Uniform Manifold Approximation and Projection (UMAP) was performed by using the RunUMAP function on the first 6 PCs to visualize cell clusters. Doublets were removed using the DoubletFinder R package (*6*). Finally, the Differential Gene Expression analysis between clusters was performed by using the non-parametric Wilcoxon Rank Sum test through the FindAllMarkers function. A gene was defined as differentially expressed (DEGs) when the Bonferroni-adjusted p-value < 0.01 and the absolute value of avg_log2FC > 0.25.

Gene Ontology enrichment analysis (GOEA) was performed on induced and inhibited genes, separately, for each Cluster, by using the DAVID Bioinformatic tool (*7*, *8*) restricting the output to Cellular Compartments (CC) terms. The threshold for statistical significance of GOEA was FDR<0.1 and Enrichment Score >1.5. The top 10 significant CCs for clusters 2 and 10 are shown in Figure 5C.

To infer TFs from single-cell data, we used the pySCENIC tool (version 0.12.1) (*9*). The pySCENIC output was integrated into the Scanpy object (*12*), and the most active regulons were identified for cells belonging to clusters 2 and 10. Subsequently, using the FIMO program (*10*) from the MEME Suite, we searched for the most active TFs obtained from the pySCENIC analysis that bound to the promoters of the differentially expressed genes in clusters 2 and 10. TF matrix for each gene was selected when the P value < 0.05.

***Data Visualization.*** Heatmaps (Figure 5C, 5E, 5F) were generated using the pheatmap package (*11*) from R on the transformed expression data and setting scale = 'row' to scale the data to Z-scores (by gene). For the UMAP figures, the Seurat object was imported to Scanpy (*12*) using Seurat's Convert function. UMAPs (Figure 5A and B) were plotted using seaborn.scatterplot (*13*) function and gene expression distribution along the UMAP (Figure 5E) was plotted using scanpy.pl.umap (*12*) function.

***Data availability.*** The 10x single-cell RNA-seq sequencing data are uploaded to GeneExpression Omnibus (GEO) under accession code GSE288395.

**Bulk RNA-seq**

***Library Preparation.*** Total mRNA was extracted from HeLa cells overexpressing CREB3L1, CREB3L1-CA, MITF or SP1 and quantified using the Qubit 4.0 fluorimetric Assay (Thermo Fisher Scientific). Libraries were prepared from 125 ng of total RNA using the NEGEDIA Digital mRNA-seq research grade sequencing service v2.0 (Next Generation Diagnostic srl) (*14*) which included library preparation, quality assessment and sequencing on a NovaSeq 6000 sequencing system using a single-end, 100 cycle strategy (Illumina Inc.).

***Bioinformatics workflow.*** The raw data were analyzed by Next Generation Diagnostic srl proprietary NEGEDIA Digital mRNA-seq pipeline (v2.0) which involves a cleaning step by quality filtering and trimming, alignment to the reference genome and counting by gene (*15*, *16*). The raw expression data were normalized, analyzed by NEGEDIA degsanalysis pipeline (v1.2.0) (*17*).

***Functional analysis on transcriptomics data*.** The threshold for the statistical significance of gene expression was FDR<0,05. To dissect which biological pathways are activated following the overexpression fo the transcription factors (MITF, CREB3L1 and SP1), Gene Ontology enrichment analysis (GOEA) was performed on induced and inhibited genes, separately, by using the DAVID Bioinformatic tool (*18*) restricting the output to Biological Process (BP), Cellular Compartments (CC) terms. The threshold for statistical significance of GOEA was FDR<0,1 and Enrichment Score ≥1.5.

***Data availability*.** The bulk RNA-seq sequencing data are uploaded to GeneExpression Omnibus (GEO) under accession code GSE299965.

**Statistical Analyses**

Statistical analyses for single cell or bulk RNA-seq data were conducted as described in the above corresponding sections. For other experiments the analyses were done using GraphPad Prism software. Unless otherwise specified, all experiments were performed at least three times. Statistical significance was determined using Student's t-test or one way ANOVA.

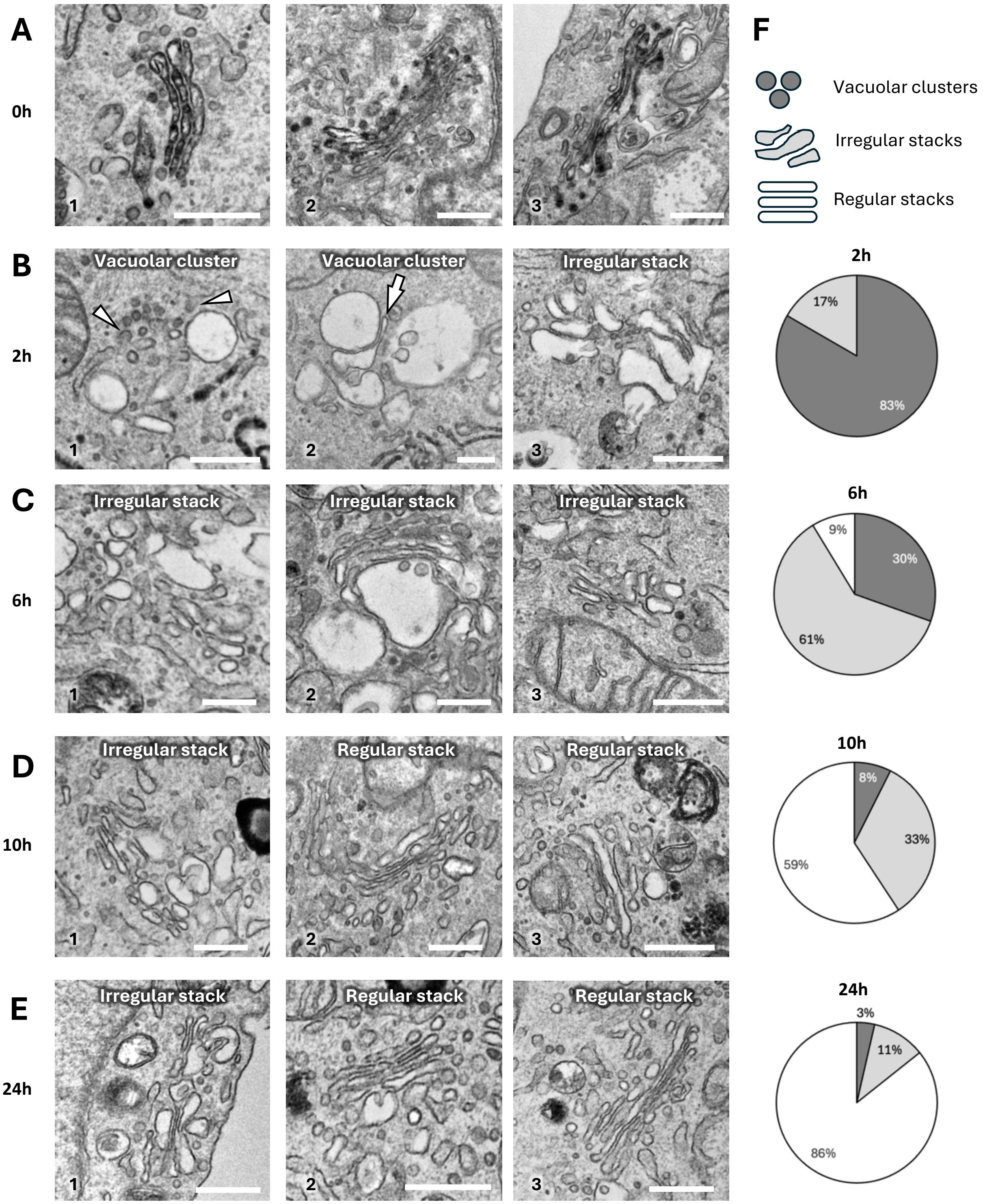


**Figure S1. Examples of Golgi recovery phenotypes at different post-inactivation time points.**
**A–E.** HeLa cells expressing ManII-HRP were incubated with DAB and $H_2O_2$ to cross-link the preexisting (old) Golgi, then fixed for EM either immediately (A) or after 2 (B), 6 (C), 10 (D), or 24 (E) hours of incubation in fresh medium. Panel A shows representative DAB deposition across the Golgi immediately after inactivation. Panels B–E show representative phenotypes of newly forming Golgi structures at each indicated time point. Arrowheads in B1 indicate an ER exit site with emerging buds and a flanking vesicular-tubular cluster. The arrow in B2 indicates a vacuolar membrane, part of which is undergoing flattening. **F.** Pie charts show the proportion of each phenotype within the population of newly forming Golgi structures at each time point. Scale bars: 320 nm (A–E).

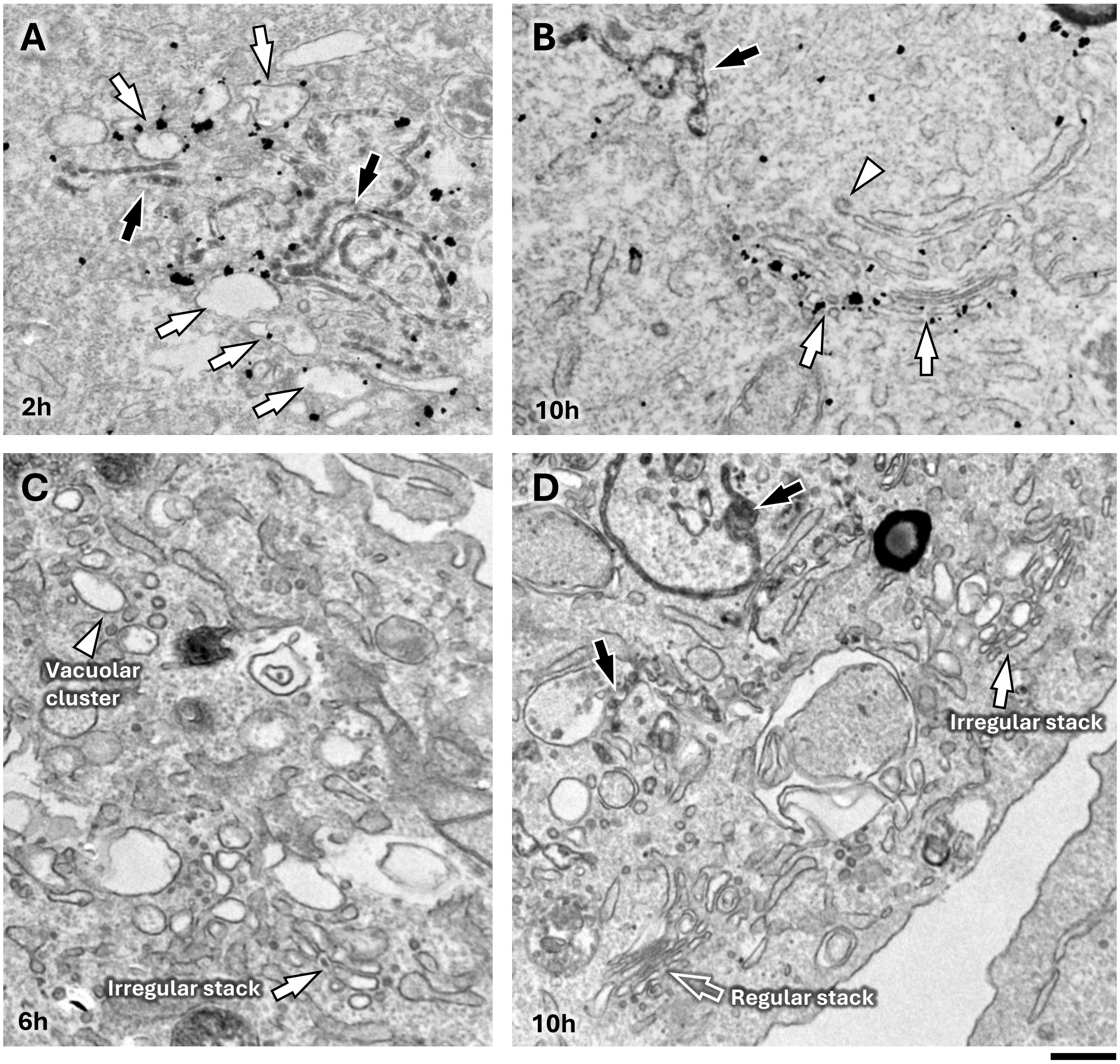


**Figure S2. Ultrastructural features of newly-forming Golgi structures.**
ManII-HRP-expressing HeLa cells were incubated with DAB and H2O2 to cross-link the preexisting (old) Golgi, fixed after incubation with fresh medium for 2 (A), 6 (C), or 10 (B, D) hours and prepared for EM. **A-B.** After fixation the cells were immuno-gold labelled for GM130. White arrows in A show vacuolar clusters decorated by gold particles associated with GM130. White arrows in B indicate GM130-positive cis-side of the Golgi, while white arrowhead shows clathrin/coated bud in the trans-portion of the stack. Black arrows in A and B indicate remnants of the old DAB-positive Golgi. **C-D**. Representative images show that the same cell may contain newly forming Golgi units with different morphologies: vacuolar cluster and irregular stack (C), or irregular and regular stack (D). Scale bar: 320 nm (A-D)

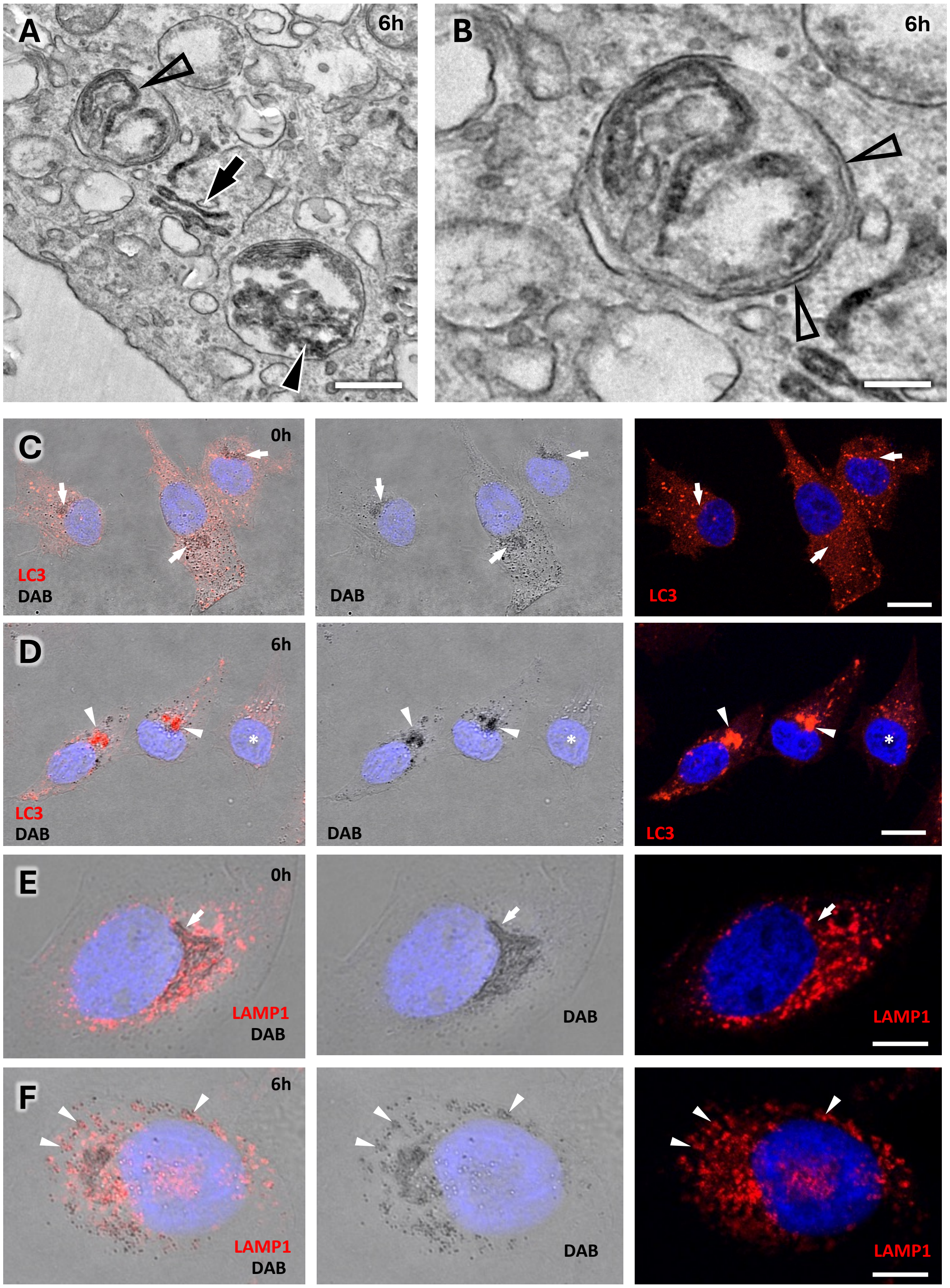


**Figure S3. Inactivated Golgi is sequestered by autophagy.**
Golgi was inactivated with DAB and H2O2 in ManII-HRP expressing HeLa cells, which were fixed immediately (C, E) or after incubation in fresh medium for 6h (A, B, D, F). **A.** Representative EM image of the cell 6h postactivation shows DAB-positive remnants of inactivated Golgi in the cytosol (black arrow) and inside autophagosome (empty arrowhead) or lysosome (black arrowhead). **B.** The image corresponds to boxed area in panel A. Arrows indicate double membrane of an autophagosome, which contains cisternae of inactivated Golgi inside. **C, D.** Cells with inactivated Golgi were labelled for LC3. DAB-positive areas of the Golgi (C, arrows) did not contain LC3 immediately after inactivation, while 6h after inactivation LC3 concentrated at the remnants of the DAB-positive old Golgi (D, arrowheads). Cell without inactivated Golgi (D, asterisk) did not exhibit similar perinuclear LC3 concentration. **E, F.** Cells with inactivated Golgi were labelled for LAMP1. Arrow in E shows inactivated Golgi in the perinuclear area. Arrowheads in F indicate DAB-positive fragments of old Golgi coinciding with LAMP1-positive spots. Scale bars: 510 nm (A), 190 nm (B), 18μm (C, D), 8μM (E, F).

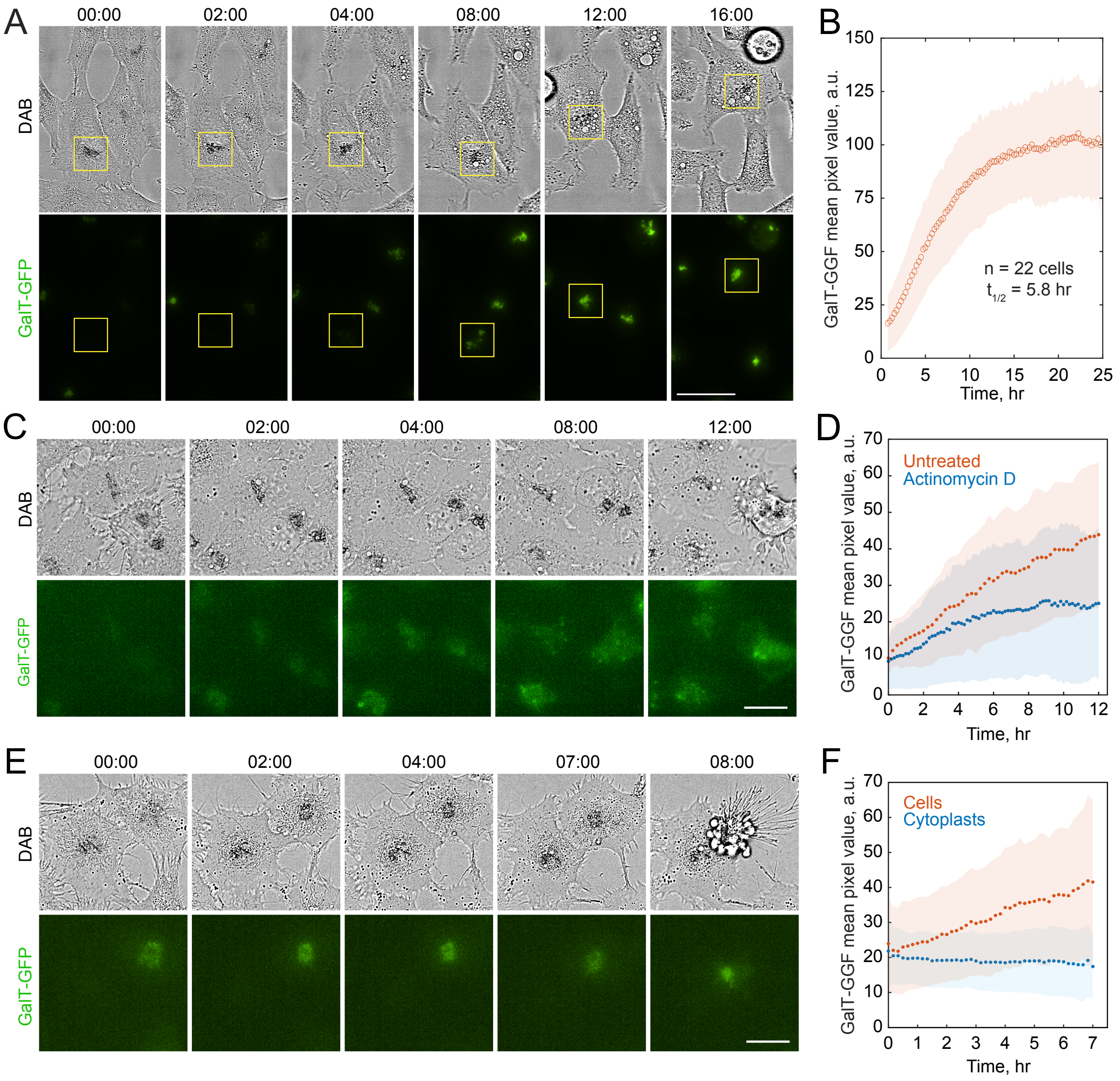


**Figure S4. Live cell imaging of Golgi recovery**
**A.** Time frames extracted from a time-lapse sequence showing Golgi recovery in *ManII-HRP* HeLa cells expressing GalT-GFP. Immediately after incubation with DAB and $H_2O_2$ (time 0), the cells display a DAB signal and no GalT-GFP fluorescence, indicating inactivation of the Golgi. The cells were then incubated with fresh medium to allow Golgi reconstitution. The square outlines the perinuclear area of the cell, where new GalT-GFP–positive Golgi structures gradually forms, while the old DAB-positive Golgi undergoes fragmentation (see Movie 2). **B.** Quantification of GalT-GFP signal over time. The fluorescence intensity reaches a plateau approximately 10–12 hours post-inactivation. **C, D.** Cells treated with DAB as in panel A were then incubated with medium containing 1 µg/ml Actinomycin D. Time-lapse imaging (C; see Movie 7) shows a significant impairment in new Golgi formation, as confirmed by quantification of GalT-GFP fluorescence (D). **E, F.** Cytoplasts were obtained by enucleation of cells expressing ManII-HRP and GalT-GFP (see Methods). After Golgi inactivation with DAB, cytoplasts fail to recover GalT-GFP signal (E; see Movie 8), as confirmed by quantification of GalT-GFP fluorescence (F). **Scale bars:** 40 µm (A), 20 µm (C, E). Time shown as HH:MM. In E and E the LUT is set to the global min and max of the signal within the sequence to reveal minimal residual GalT-GFP fluorescence remaining after quenching by DAB.

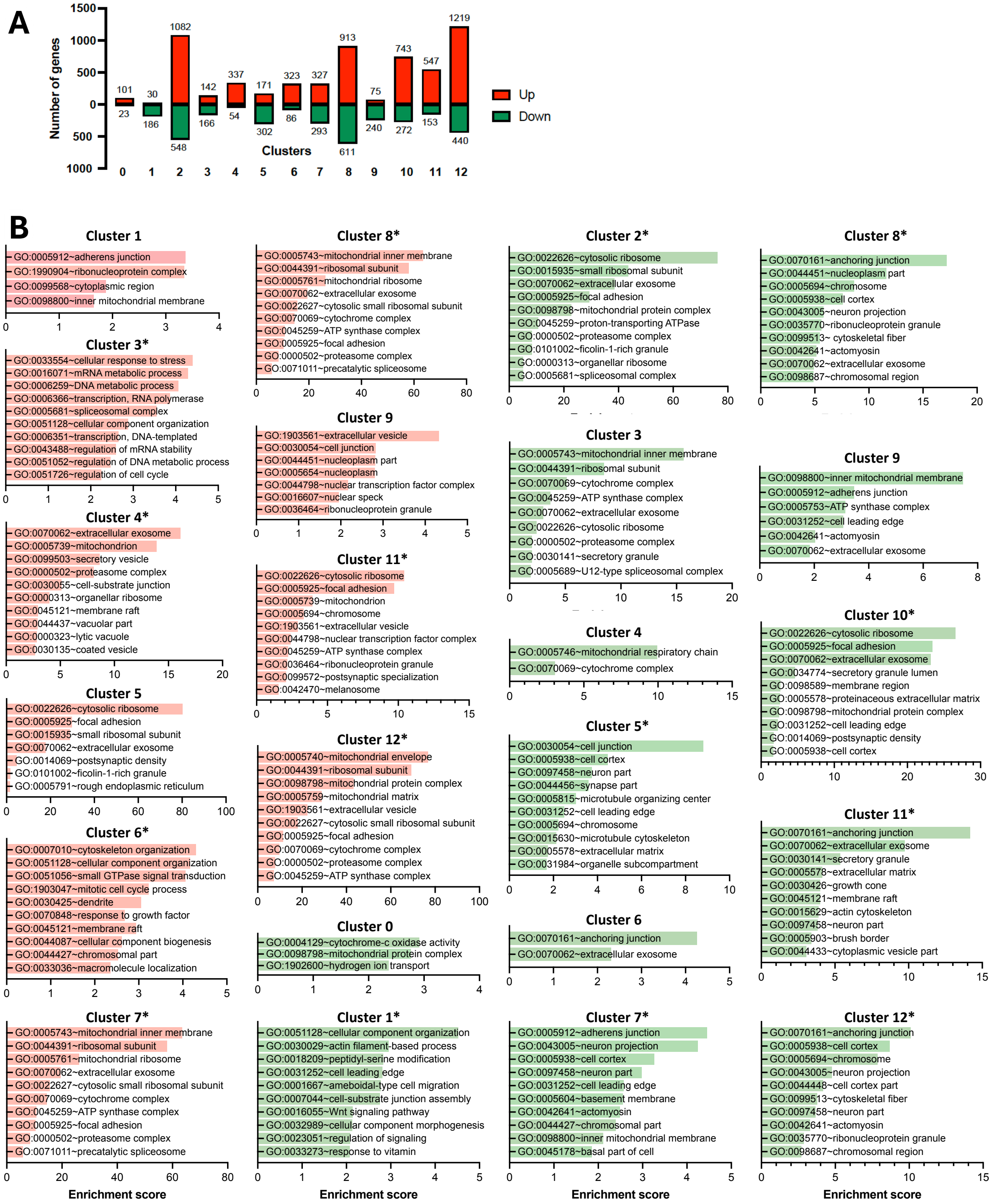


**Sup Figure 5. Gene ontology (GO) enrichment analysis of differently expressed genes (DEGs) in each cluster of scRNA-sec dataset.**

DEGs were identified (as described in Material and Methods) in each cluster of cells (see Trans Fig 1A), which were subjected to Golgi inactivation and subsequent RNAseq analysis. **A.** The graph shows number of DEGs for each cluster. **B.** Differently expressed genes in each cluster were subjected to GO enrichment analysis. The graphs show enriched annotation terms for up- (red) or down-regulated (green) transcripts in each cluster. Enrichment scores for upregulated genes in clusters 2 and 10 are shown in main Fig 5C. No significant enrichments were detected for upregulated genes in cluster 0. Asterisk indicate clusters for which only top 10 terms are shown.

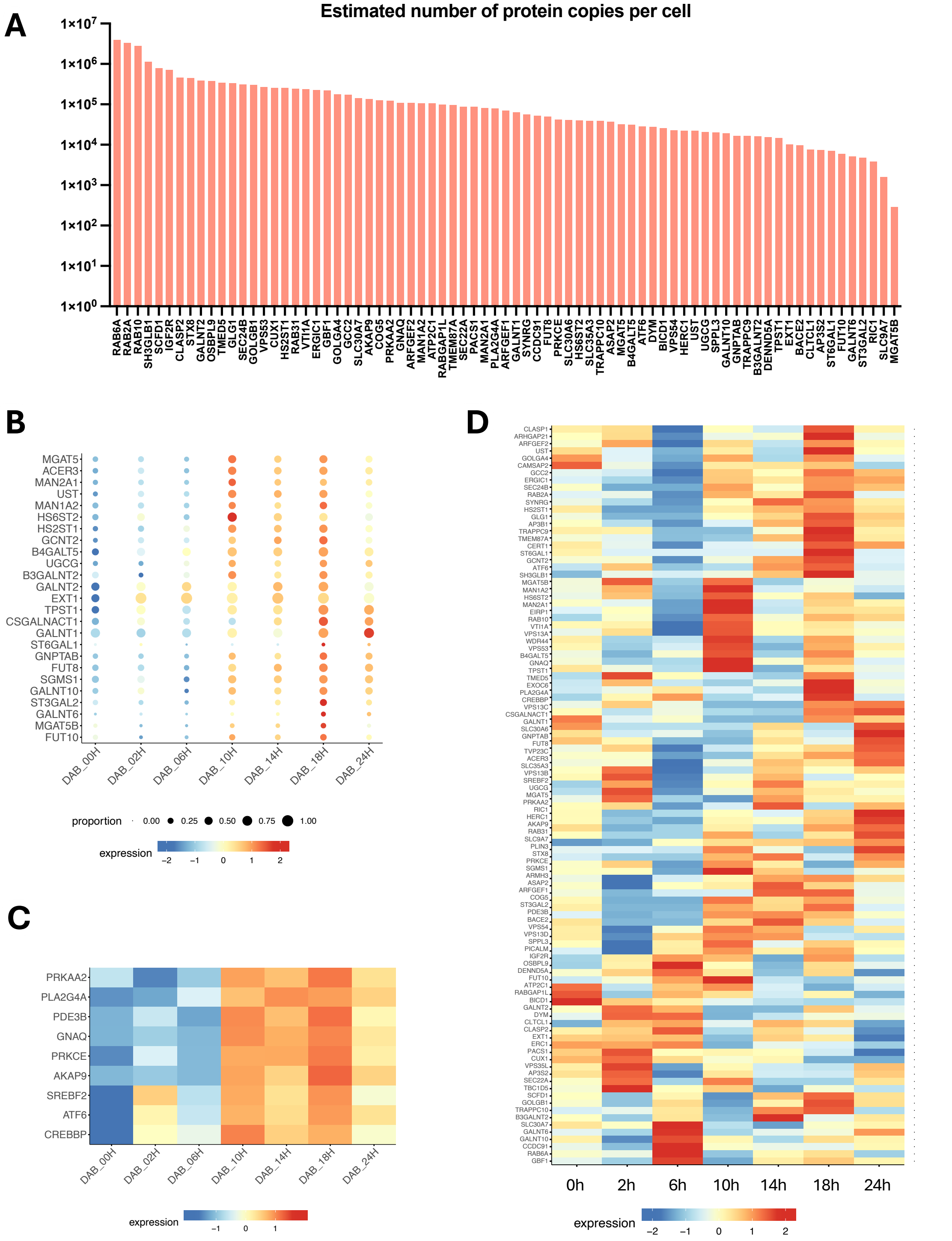


**Figure S6. Specific features of activated Golgi genes.**
**A.** The estimated numbers of copies of Golgi proteins per cell were recovered from published proteomics data (Fasimoye et al., 2023) and plotted for corresponding genes, whose activation was observed during biogenesis of the new Golgi. **B.** The bubble plot shows a subset of the Golgi genes (encoding glycosylation enzymes). Both the expression levels and number of cells expressing these genes increase with after inactivation of the old Golgi, especially at 10h timepoint. **C.** The heat map shows expression dynamics of genes encoding Golgi-associated signaling proteins. Transcription factor associated genes ATF6, CREBBP, and SREBF2 exhibit earlier transactivation and are clustered together. **D.** The expression of the Golgi genes (same as in Fig. 6C) was evaluated at different time points after incubation with DAB in cells without detectable ManII-HRP expression. Majority of the genes exhibit fluctuating expression without specific activation time point.

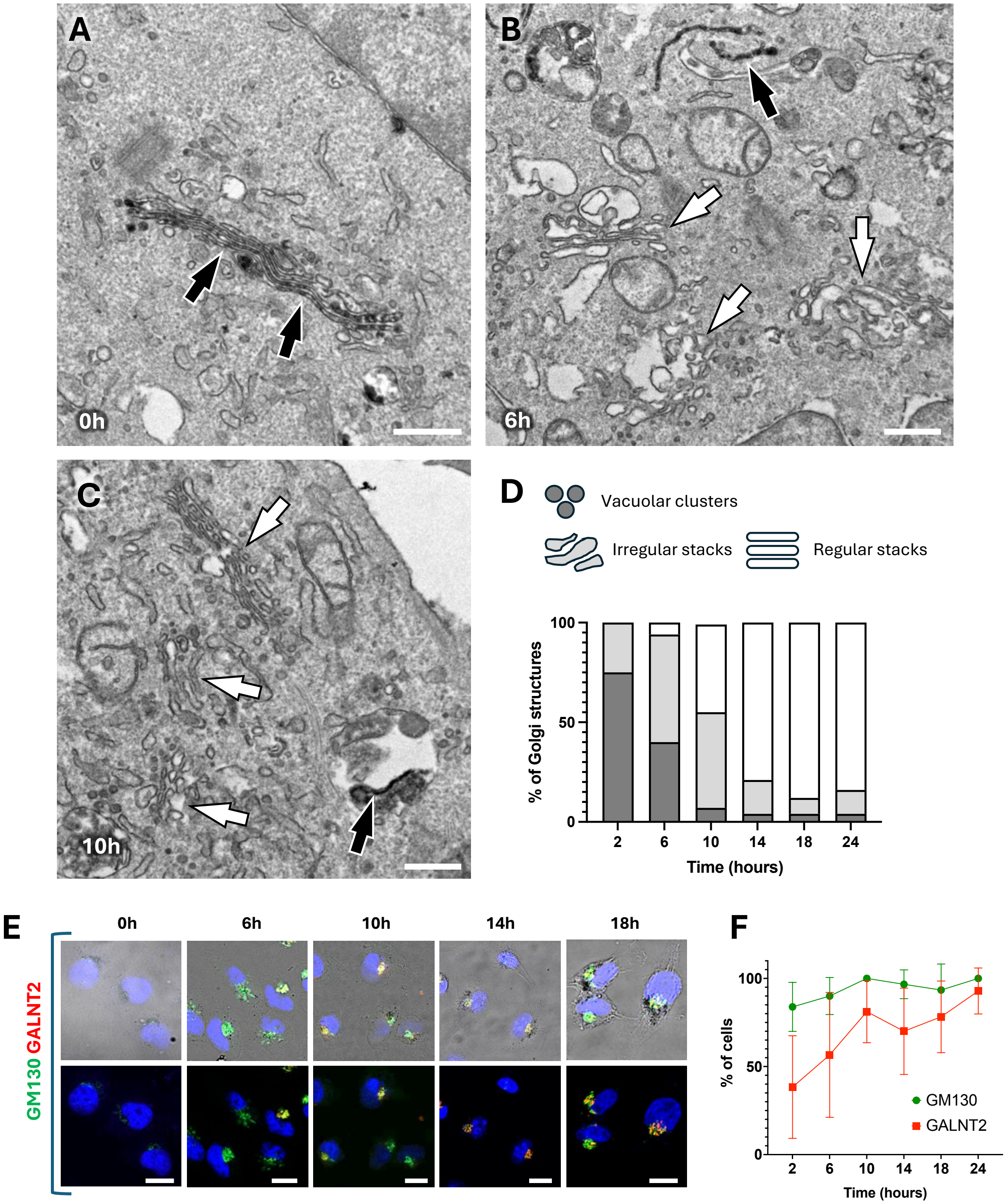


**Figure S7. Golgi biogenesis in cells processed in parallel with scRNA-seq experiment.**
ManII-HRP-expressing HeLa cells were incubated with DAB and $H_2O_2$ to cross-link the preexisting (old) Golgi. The cells were collected immediately after Golgi inactivation or at 2, 6, 10, 14, 18 and 24h intervals of incubations with fresh medium. From each time point the collected cells were processes for scRNA-seq (see figures 5 and 6), EM (A-D) and IF (E, F) analyses. **A-D**. EM images (A-C) and quantification (D) show that substantial proportion of newly formed Golgi units acquire regular stack architecture starting from 10h post-inactivation. Black and white arrows indicate old and new Golgi respectively. **E, F.** Confocal images (E) and quantification (F) show that majority of cells recover GALNT2 in the newly forming Golgi units at 10h interval after inactivation of the old Golgi. Scale bars: 360 nm (A-C), 17 µm (E).

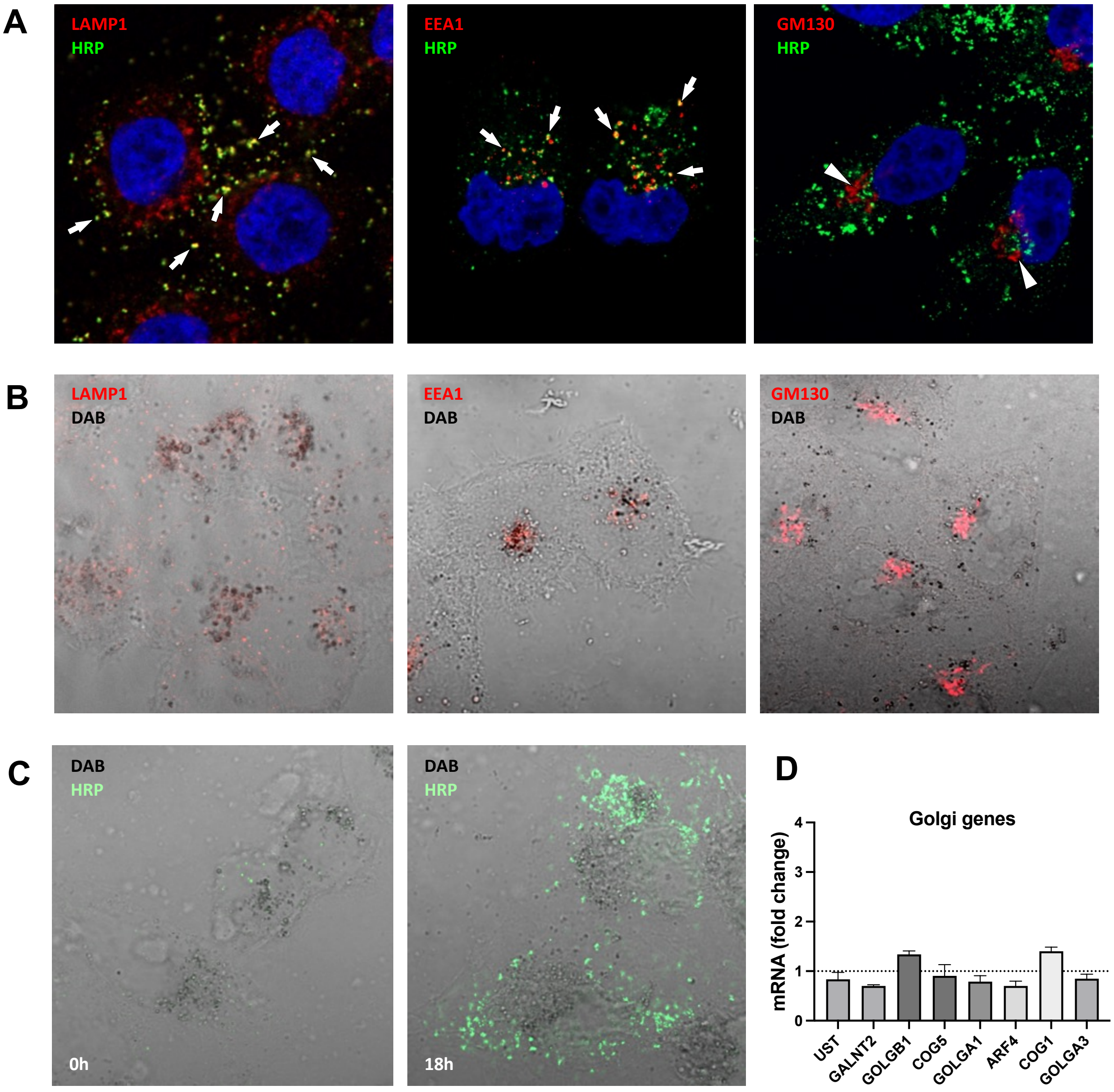


**Figure S8. Endosome inactivation does not induce Golgi gene expression.**

**A.** HeLa cells were incubated with HRP for 30 min, fixed, and stained to examine colocalization of internalized HRP with markers of late endosomes (LAMP1), early endosomes (EEA1), or the Golgi apparatus (GM130). Arrows indicate numerous EEA1- or LAMP1-positive structures containing HRP. Arrowheads mark Golgi structures that lack any HRP signal. **B.** Cells were loaded with HRP as in panel A and subsequently treated with DAB and $H_2O_2$ to inactivate HRP-loaded endosomes. The panels show substantial overlap of DAB with LAMP1 or EEA1 signals, which were also partially quenched, but not with GM130, indicating that the Golgi was not targeted by inactivation. **C.** Endosomes were inactivated as described in panel B. Cells were then incubated with HRP again for 30 min either immediately (0 h) or 18 h later to assess the efficiency of HRP endocytosis. At 0 h, very little HRP uptake was observed, confirming that endosomes were functionally disabled immediately after inactivation. In contrast, numerous HRP-positive endocytic structures were detected at 18 h post-inactivation, indicating efficient regeneration of functional endosomal organelles. **D.** To determine whether inactivation and subsequent de novo biogenesis of endosomes induces Golgi gene expression, HeLa cells were collected immediately after endosome inactivation (0 h) and at 18 h, when new endosomes had formed. This 18 h time point corresponds to the peak of Golgi gene expression during de novo Golgi biogenesis. The expression of Golgi genes was assessed by qRT-PCR, as HRP uptake and subsequent endosome inactivation were homogeneous across the cell population. The graph shows mRNA fold changes for each Golgi gene at 18 h post-inactivation relative to the 0 h time point (dashed line). No significant changes in the expression of the tested genes were detected.

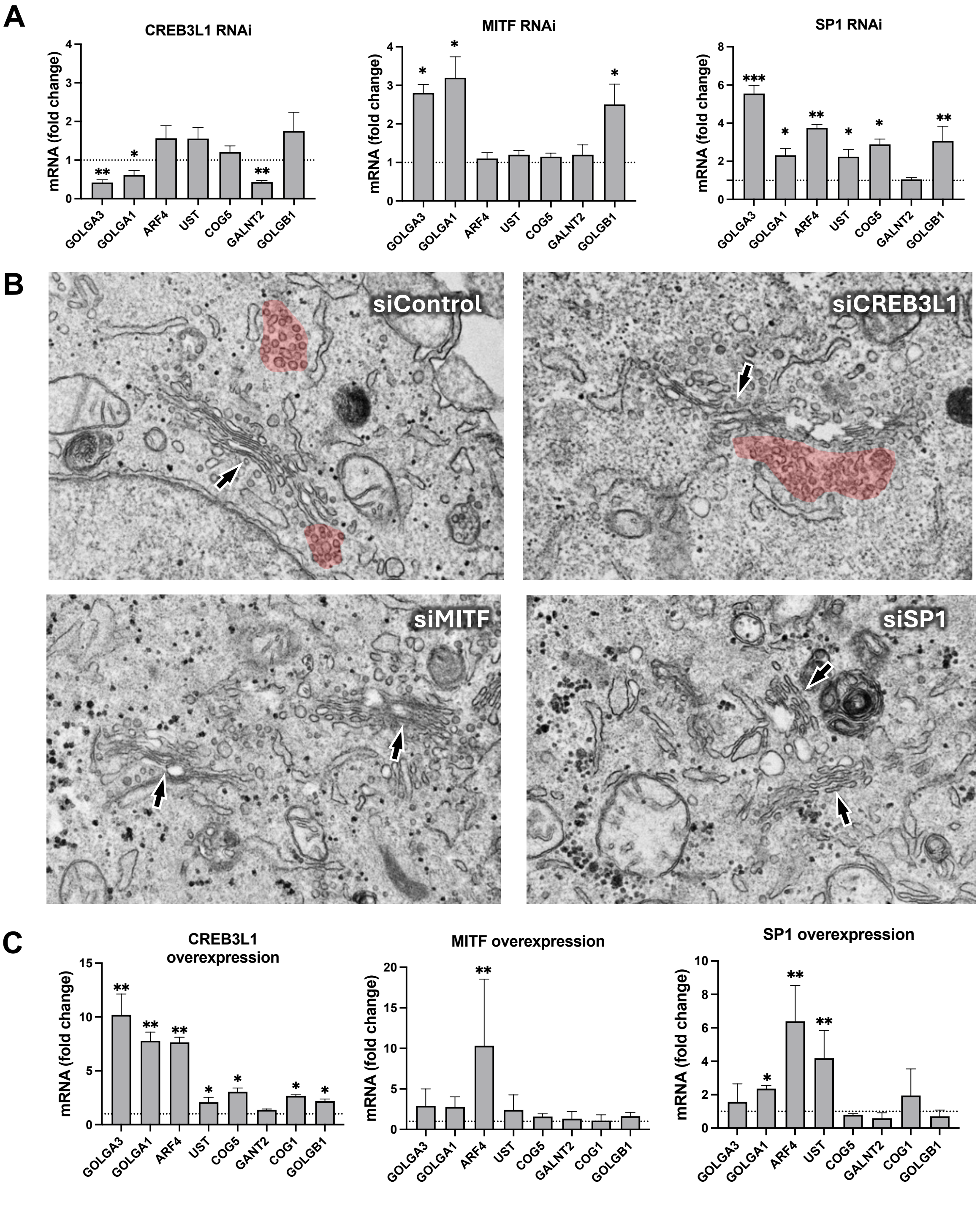


**Figure S9. Impact of silencing and overexpression of TF candidates.**

**A.** CREB3L1, MITF, or SP1 were silenced in HeLa cells, and changes in Golgi gene expression were evaluated by qRT-PCR relative to control cells (dashed line). Statistical significance: ***p<0.001, **p<0.01, *p<0.05 (one-way ANOVA; n=3 experiments). **B.** Golgi ultrastructure was assessed by electron microscopy (EM) in silenced cells. Arrows indicate Golgi stacks. Red highlights mark pre-Golgi vesicular-tubular clusters (VTCs), indicating significant VTC expansion in CREB3L1-silenced cells. **C.** CREB3L1, MITF, or SP1 were overexpressed in HeLa cells, and changes in Golgi gene expression were evaluated by qRT-PCR relative to control cells (dashed line). Statistical significance: **p<0.01, *p<0.05 (one-way ANOVA; n=3 experiments).

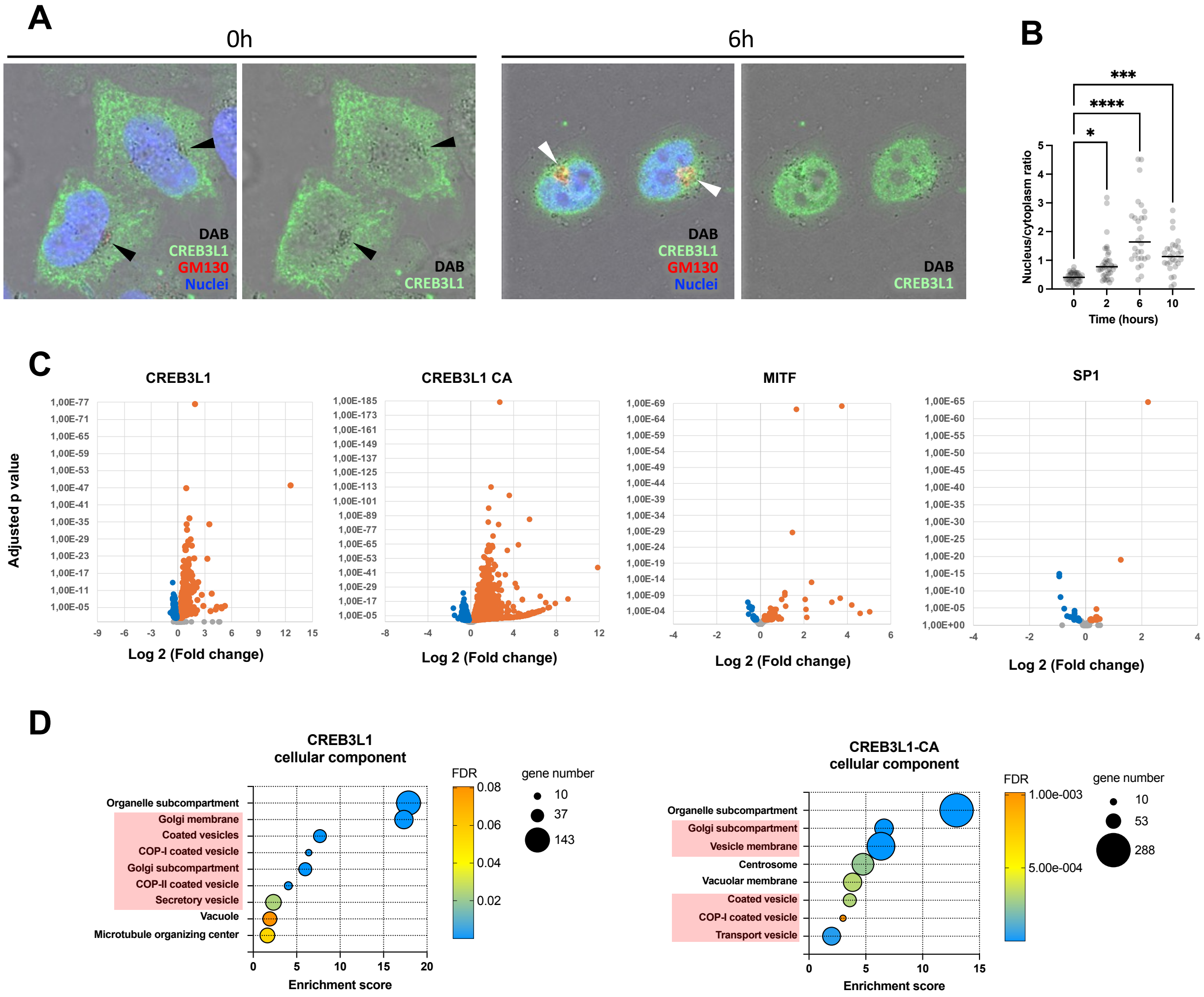


**Figure S10. CREB3L1 activation and its impact on Golgi-related genes**

**A, B.** ManII-HRP cells were transfected with CREB3L1 and subjected to Golgi inactivation. In panel A, black arrows indicate DAB-labeled, crosslinked Golgi. After inactivation, cells were either fixed immediately (0 h) or incubated in fresh medium for various time points (2, 6, or 10 h) to allow Golgi reconstitution. Cells were then fixed and immunostained for CREB3L1, GM130, and nuclei. A prominent translocation of CREB3L1 to the nucleus was observed at 6 h post-inactivation. White arrows in panel A indicate newly forming Golgi structures. Quantification in panel B shows an increase in the nuclear-to-cytoplasmic ratio of CREB3L1 signal, peaking at 6 h (****$p < 0.0001$, ***$p < 0.001$, *$p < 0.05$; one-way ANOVA; $n = 30$ cells). **C.** HeLa cells were transfected with CREB3L1, its constitutively active form (CREB3L1-CA), MITF, or SP1, followed by bulk RNA-seq analysis. Volcano plots illustrate the overall transcriptional impact of each transcription factor, with significantly upregulated and downregulated genes highlighted in orange and green, respectively. CREB3L1 and CREB3L1-CA induced significantly broader transcriptomic changes compared to the other factors. **D.** Gene Ontology enrichment analysis of upregulated transcripts in CREB3L1- and CREB3L1-CA-expressing cells revealed significant enrichment in Golgi-related functional annotation terms (highlighted in pink).

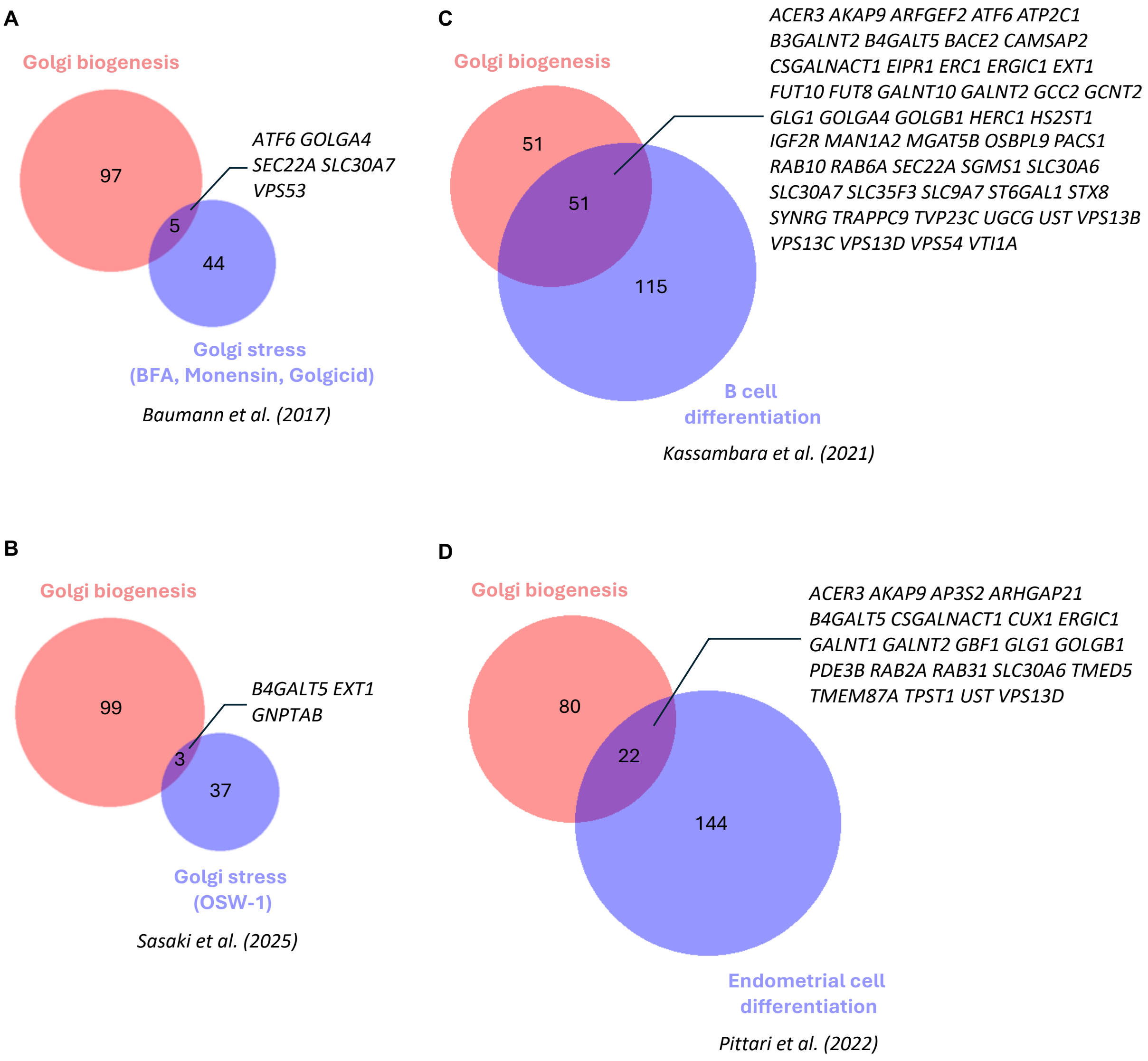


**Figure S11. Comparison of Golgi gene sets activated during Golgi bogenesis, Golgi stress, or cell differentiation.**
Venn diagrams show the overlap between set of Golgi genes activated during Golgi biogenesis (this study) and Golgi gene sets reported to be induced by Golgi stress (A, B) or during cell differentiation (C, D). For each panel, the number of overlapping and non-overlapping genes is indicated. Common gene names are listed, and the source of each compared gene set is referenced.

**Movie S1. Rapid DAB accumulation in HeLa cells following Golgi cross-linking.**
The movie begins immediately after HeLa cells expressing ManII-HRP and GalT-GFP were incubated with diaminobenzidine (DAB) and hydrogen peroxide ($H_2O_2$) to cross-link the pre-existing Golgi. DAB accumulation within the Golgi region is observed within 30 seconds and is accompanied by quenching of GalT-GFP fluorescence.

**Movie S2. Golgi recovery after inactivation.**
The movie begins immediately after ManII-HRP HeLa cells expressing GalT-GFP were incubated with DAB and $H_2O_2$, resulting in quenching of the GalT-GFP signal in the pre-existing Golgi. Fluorescence reappears in the newly forming Golgi a few hours after inactivation and reaches a plateau around 10–12 hours.

**Movie S3. Trafficking of GPI-RUSH in control cells.**
This time-lapse sequence shows trafficking of GPI-RUSH released from the ER in control ManII-HRP HeLa cells upon biotin addition. Bright-field images at the beginning confirm the absence of DAB deposition in the Golgi. After ER release, GPI-RUSH rapidly traffics to the Golgi and subsequently to the cell surface.

**Movie S4. Trafficking of GPI-RUSH during Golgi recovery after inactivation.**
This time-lapse sequence shows trafficking of GPI-RUSH released from the ER in ManII-HRP HeLa cells treated with DAB to cross-link the Golgi. Biotin addition triggers GPI-RUSH trafficking immediately after Golgi inactivation. While ER-to-Golgi transport occurs at speeds comparable to control cells, subsequent export from the Golgi and delivery to the plasma membrane is markedly slowed in DAB-treated cells.

**Movie S5. Trafficking of M6PR-RUSH in control cells.**
This movie shows trafficking of M6PR-RUSH released from the ER in control ManII-HRP HeLa cells. Following ER release, M6PR-RUSH (red) rapidly traffics to the Golgi, highlighted by GalT-GFP (green), and then moves from the Golgi to scattered post-Golgi endosomal structures.

**Movie S6. Trafficking of M6PR-RUSH during Golgi recovery after inactivation.**
This movie shows trafficking of M6PR-RUSH released from the ER in ManII-HRP HeLa cells treated with DAB to cross-link the Golgi. Biotin addition initiates M6PR-RUSH trafficking immediately after Golgi inactivation (DAB deposition is visible in the bright-field panel). After ER exit, M6PR-RUSH (red) gradually accumulates within the newly forming Golgi (green); however, its export to endosomal compartments is substantially delayed compared to control cells.

**Movie S7. Golgi recovery is impaired upon transcription inhibition.**
The Golgi complex was inactivated in ManII-HRP cells expressing GalT-GFP, followed by treatment with the transcription inhibitor Actinomycin D (Act-D). These cells failed to recover the Golgi, as indicated by the absence of recovery of the compact GalT-GFP fluorescent signal in the perinuclear region. Note: the LUT was set to the global minimum and maximum of the signal within the sequence to visualize minimal residual GalT-GFP fluorescence remaining after DAB quenching.

**Movie S8. Golgi recovery is impaired upon cell enucleation.**
ManII-HRP cells expressing GalT-GFP were enucleated prior to inactivation of the pre-existing Golgi. Nucleus-free cytoplasts, which lack transcriptional activity, failed to rebuild the Golgi apparatus. Note: the LUT was set to the global minimum and maximum of the signal within the sequence to visualize minimal residual GalT-GFP fluorescence after DAB quenching. A residual GalT-GFP signal is visible in one cytoplast (on the right), but it does not increase over time.

**Data S1. Set of Golgi genes activated during Golgi biogenesis.**

**Data S2. Gene Ontology enrichment analysis (annotation clusters) of differently expressed genes in CREB3L1 overexpressing cells.**

**Data S3. Gene Ontology enrichment analysis (annotation clusters) of differently expressed genes in CREB3L1-CA overexpressing cells.**

**Data S4. Gene Ontology enrichment analysis of differently expressed genes in MITF overexpressing cells.**

**Data S5. Gene Ontology enrichment analysis of differently expressed genes in SP1 overexpressing cells.**

**Data S6. List of the siRNAs used in the study.**

**Data S7. List of primers used in the study.**
